# Network structure influences the strength of learned neural representations

**DOI:** 10.1101/2023.01.23.525254

**Authors:** Ari E. Kahn, Karol Szymula, Sophie Loman, Edda B. Haggerty, Nathaniel Nyema, Geoffrey K. Aguirre, Dani S. Bassett

## Abstract

Human experience is built upon sequences of discrete events. From those sequences, humans build impressively accurate models of their world. This process has been referred to as graph learning, a form of structure learning in which the mental model encodes the graph of event-to-event transition probabilities [1], [2], typically in medial temporal cortex [3]–[6]. Recent evidence suggests that some network structures are easier to learn than others [7]–[9], but the neural properties of this effect remain unknown. Here we use fMRI to show that the network structure of a temporal sequence of stimuli influences the fidelity with which those stimuli are represented in the brain. Healthy adult human participants learned a set of stimulus-motor associations following one of two graph structures. The design of our experiment allowed us to separate regional sensitivity to the structural, stimulus, and motor response components of the task. As expected, whereas the motor response could be decoded from neural representations in postcentral gyrus, the shape of the stimulus could be decoded from lateral occipital cortex. The structure of the graph impacted the nature of neural representations: when the graph was modular as opposed to lattice-like, BOLD representations in visual areas better predicted trial identity in a held-out run and displayed higher intrinsic dimensionality. Our results demonstrate that even over relatively short timescales, graph structure determines the fidelity of event representations as well as the dimensionality of the space in which those representations are encoded. More broadly, our study shows that network context influences the strength of learned neural representations, motivating future work in the design, optimization, and adaptation of network contexts for distinct types of learning over different timescales.

## Introduction

Human experience is composed of events, words, and locations that occur in sequences, following predictable patterns. For example, heading to work each day, your route is formed by a series of intersections, as could be conveyed in turn-by-turn directions. Over time, you construct a mental map of not just the intersections but also the route. Likewise, when speaking to a colleague, you predict syllables and words based on your knowledge of the language and scholarly field, as evident in your ability to fill in gaps in the sounds and words that reach you due to, for example, background noise. Even when watching a film, our minds synthesize a temporal sequence of scenes into a multifaceted set of interactions between characters and events. The formation of such mental representations depends upon our ability to detect hidden interactions and to transform them into a predictive model.

Building such statistical models is central to human cognition [9], [10], because it allows us to accurately predict and respond to future events. One of the most prominent cues to underlying structure is the set of transition statistics between elements, such as how frequently one word precedes another in a lexicon. This cue has been extensively studied in the field of statistical learning [11], and used to demonstrate that syllable transitions can help demarcate word boundaries as infants learn language [12], [13]. Such statistical learning occurs in both infants (see [14] for an overview) and adults [15] and is thought to be a fundamental, domain-general learning mechanism because it arises from various sorts of sequences—from tones [16] and shapes [17] to motor actions [18]. Human learners likewise exhibit sensitivity to statistics between non-adjacent elements in a sequence [19], [20]. Despite these strands of evidence, a comprehensive understanding of such non-adjacent statistics is challenging due to the number of possible statistical relations on which human learners might rely.

To study the role of non-adjacent probabilities in statistical learning, one particularly promising approach explicitly models a collection of events and the transitions between them as a graph [21], [22]. Here nodes in the graph represent events and edges represent possible transitions between any two events. By formally mapping transition probabilities to a graph, one can study how the graph’s topological properties influence human learning. Practically, an experimenter can construct a sequence of events by a walk on the graph, and then assess the participant’s ability to predict upcoming items by measuring their response time to a cover task. Studies of this ilk have demonstrated that topological features impacting learnability include the type of walk [23] and whether the graph contains densely interconnected subsets or *modules* of nodes [7], a property characteristic of many real-world networks shown to efficiently convey information [9]. Interestingly, it remains unknown what neural processes might explain the relation between graph topology and behavior.

In contrast to prior work exploring neural representations of items within the same graph [24], [25], here we designed a study to ask whether neural representations of stimuli systematically differed between different graph structures. In particular, given the ability of modular structure to efficiently convey information [9], we hypothesized that the presence of modular structure would lead to robust neural representations, whereas the lack of modular structure would lead to weak neural representations. As a comparison structure, we chose a ring lattice graph, which allowed us to control for local degree while eliminating modularity. To test our hypothesis, we measured BOLD fMRI activity while participants learned to predict and respond to sequences of abstract shapes. We found that modular structure led to improved discriminability between the neural activity patterns of stimuli, as well as higher intrinsic dimensionality, when compared to the ring lattice. These results demonstrate that neural representations differ systematically in response to different graph structures. Moreover, the work suggests that robust behavior may depend upon our ability to extract organizational patterns such as graph modularity. More broadly, our findings underscore the need to characterize how networks of information shape human learning.

## Results

Thirty-four participants (11 male and 23 female) were recruited from the general population of Philadelphia, Pennsylvania USA. All participants gave written informed consent and the Institutional Review Board at the University of Pennsylvania approved all procedures. All participants were between the ages of 18 and 34 years (M=26.5 years; SD=4.73), were right-handed, and met standard MRI safety criteria. Two of the thirty-four participants were excluded because they failed to learn the shape-motor associations (one failed to complete the pre-training; one answered at-chance during the second session), and one additional participant was excluded because we did not accurately record their behavior due to technical difficulties. Hence, all reported analyses used data from thirty-one participants.

### Task

Each participant completed 2 sessions of a visuo-motor response task in an MRI scanner while BOLD activity was recorded. The visuo-motor task consisted of a set of shapes, a set of motor responses, and a graph which both served to map the shapes to responses and to generate trial orderings. Each node in the graph was associated with a shape and a motor response. For the former, we designed a set of 15 abstract shapes (Fig. 1a, left) to be visually discriminable from one another and to evoke activity in the lateral occipital cortex. For the latter, we chose the set of all 15 one- or two-button motor responses on a five-button response pad (Fig. 1a, right). We assigned participants to one of two conditions: a modular graph condition (*n* = 16) or a ring lattice condition (*n* = 15) (Fig. 1b, left). Each trial corresponded to a node on the graph, and trial order was determined by a walk between connected nodes. A three-way association between node, shape, and motor response was generated, at random, for each participant: each of the 15 graph nodes was assigned a unique shape and a unique motor response, both chosen at random from the set of shapes and the set of motor responses, respectively (Fig. 1b, right). The mappings of shapes and motor responses were independent of one another, and randomized between participants, allowing us to separate the effects of shape, motor response, and graph.

**Figure 1:**
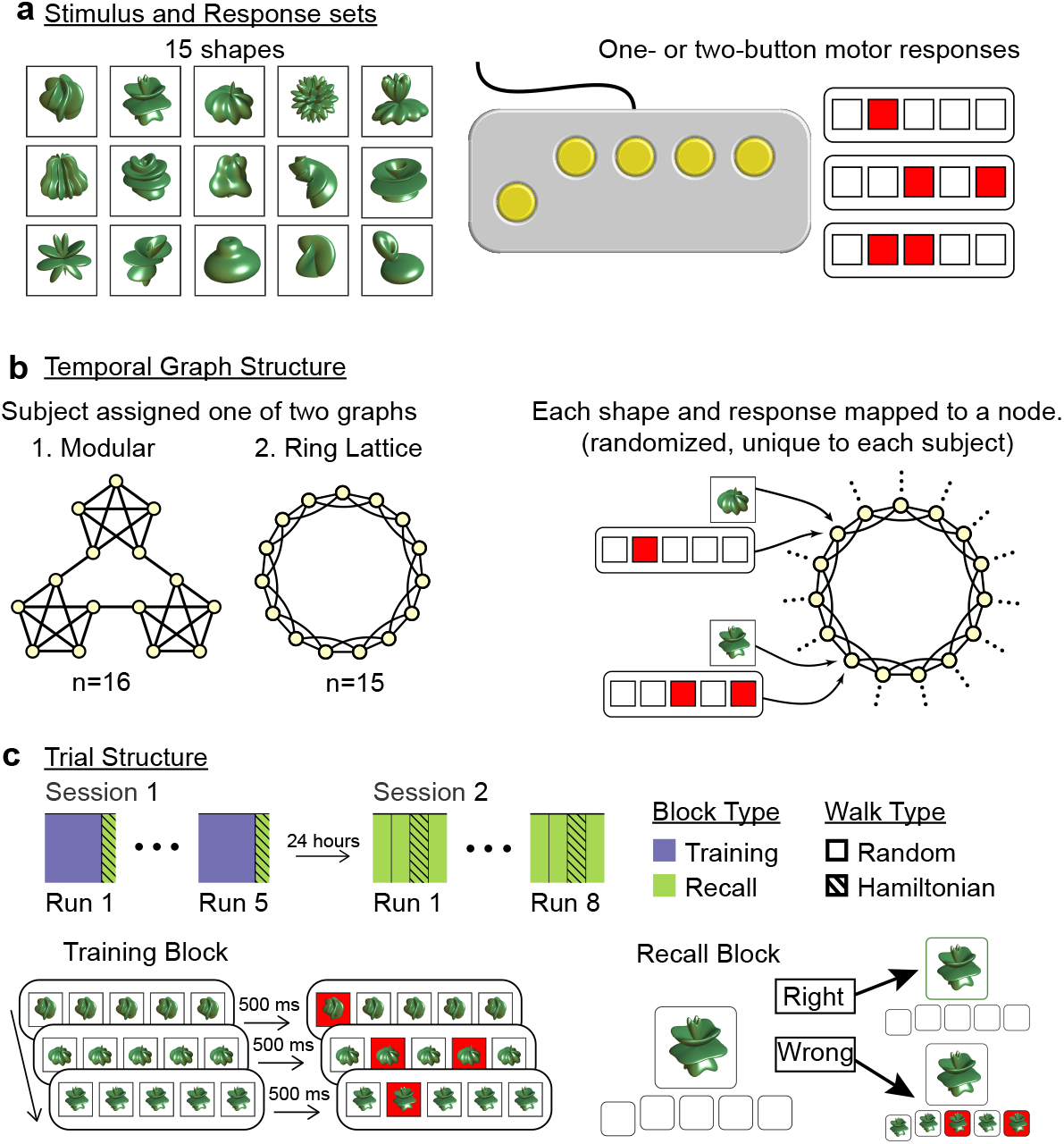
The structure of the graph learning experiment, which incorporated visual stimuli and motor responses on two graph topologies. (**a**) Participants were trained and tested on a set of 15 shapes (*left*) and 15 possible one- or two-button combinations on a response pad (*right*). (**b**) The order of those trials, and how the shapes and motor responses related to one another, varied between participants and across the two graph conditions. Each participant was assigned to one of two graph structures: modular or ring lattice (*left*). Then, each of the 15 shapes and motor responses was mapped to one of the 15 nodes in the assigned graph (*right*). To control for differences among shapes and responses, each mapping was random and unique to each participant. (**c**) Session one (*top left*) comprised five runs of a training block followed by a recall block. Session two (*top right*) comprised eight runs of four recall blocks each. Each block was composed of a series of trials, following a random or Hamiltonian walk on the assigned graph. In training blocks (*bottom left*), participants were instructed to press the buttons indicated by the red squares. To encourage participants to learn shape mappings, the shape appeared 500 ms prior to the motor command. In recall blocks (*bottom right*), the shape was shown and participants had two seconds to respond. If they responded correctly, then the shape was outlined in green; if they response incorrectly, then the correct response was shown.

The experiment extended over two days. Session one occurred on the first day and session two occurred on the second day. Session one consisted of five task runs, and each run consisted of a training block and a recall block. Session two consisted of eight runs of four recall blocks each (Fig. 1c, top). During each training block, five squares were shown on the screen, horizontally arranged to mimic the layout of the response pad. On each trial, two redundant cues were presented: (i) a shape was shown in all five squares, and (ii) the associated motor response was indicated by highlighting one or two squares in red. Either cue alone was sufficient to pick the correct response. The shape was displayed 500 ms earlier than the motor movement in order to encourage participants to learn the visuo-motor pairings. Participants completed 300 trials at their own pace: each trial would not advance until the correct motor response was produced. Once produced, the shape for the next trial would immediately appear, and 500 ms later the square(s) corresponding to the next required motor response would be highlighted. During each recall block, the shape alone was displayed in the center of the screen, and participants were given two seconds to produce the correct motor response by memory. No motor cue was given, unless the participant answered incorrectly: in the latter case, the correct response was shown below the shape for the remainder of the two seconds (Fig. 1c, bottom).

### Behavior

We measured response accuracy to verify that participants learned the task structure and the visuo-motor associations. For participants to attain high response accuracy during the training blocks, they must simply respond to the motor commands presented to them; in contrast, for participants to attain high response accuracy during the recall blocks, they must correctly learn the shape-motor associations. In the training blocks of session one, we observed a mean accuracy of 89.1% by run five, suggesting that participants could produce the motor response with high accuracy when explicitly cued (Fig. 2a, left). In the recall blocks of session one (Fig. 2a, center), we observed a mean accuracy of 74.4% by run five (chance was 6.67%), indicating that participants could also produce the motor response with some accuracy despite the absence of any motor cue. At the beginning of session two, the recall accuracy was slightly lower (at 70.8%), but by the end of session two, the recall accuracy was 96.0%, demonstrating robust recall of motor responses (Fig. 2a, right). Collectively, these data demonstrate that participants were able to learn the motor response assigned to each visual stimuli over a two-day period.

**Figure 2:**
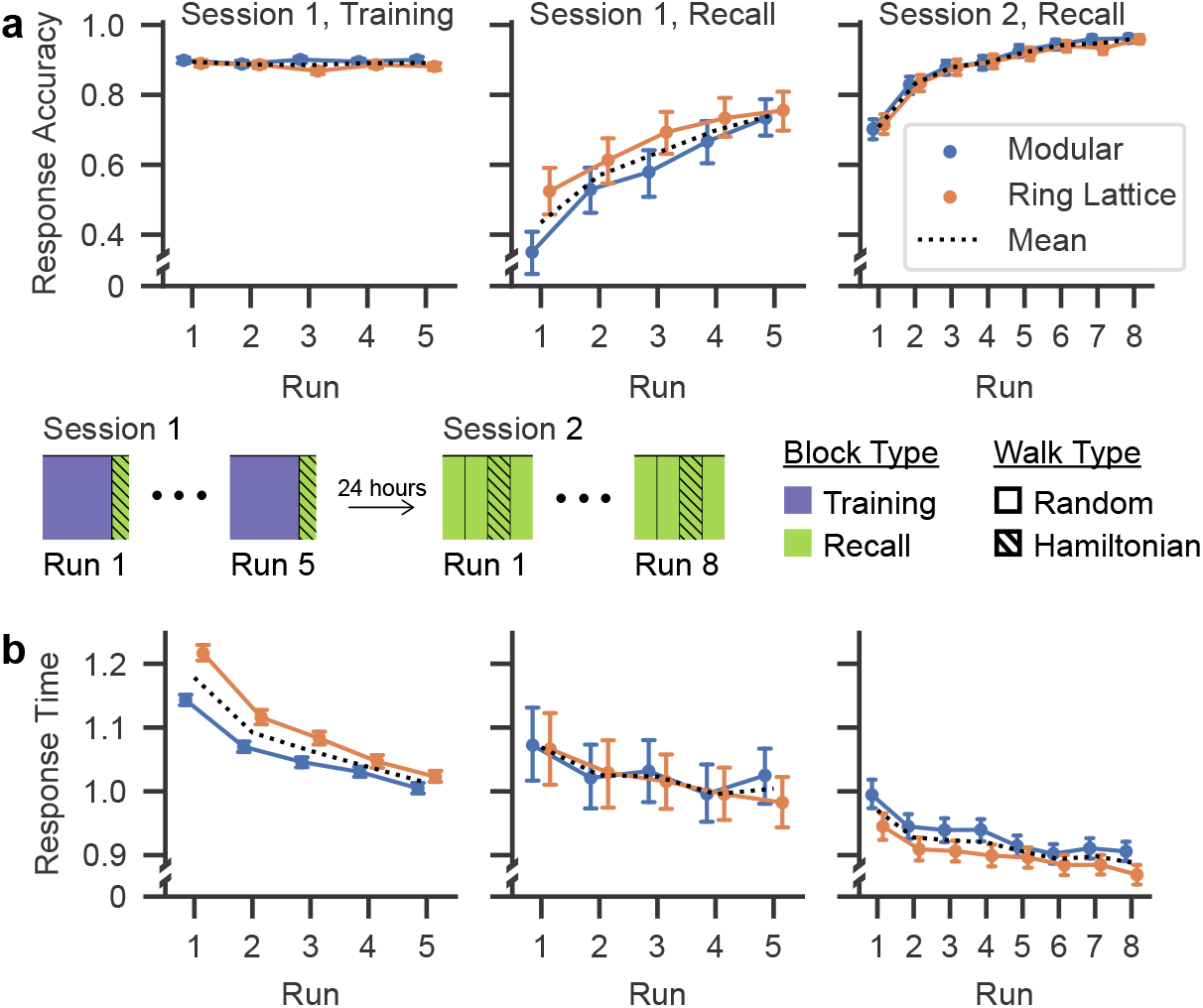
Participant behavior in the graph learning experiment. (**a**) Participant response accuracy. Markers indicate the mean values and error bars indicate the 95% confidence intervals across participant averages for each block and run. (**b**) Participant response times for correct trials. Markers indicate the mean values and error bars indicate the 95% confidence intervals across participant averages for each block and run. In both panels **a** and **b**, the blue (orange) line indicates quantities calculated from data acquired from participants assigned to the modular (ring lattice) graph condition. The black line indicates the mean across both graph conditions. In both panels, we show quantities calculated for session one training (*left*) and recall (*center*) blocks, as well as for session two recall blocks (*right*).

Whereas response accuracy allows us to assess the learning of visuo-motor associations, response time allows us to assess each participant’s expectations regarding event structure and the underlying graph that encodes that structure. If a participant is able to learn the underlying graph structure, then they can predict which shapes might be coming next (those directly connected to the current shape in the graph), and will respond more quickly. Consistent with this intuition, we found that response times steadily decreased during training, from approximately 1.18 s during run one to approximately 1.01 s during run five (Fig. 2b, left). During the recall blocks, response times depended on the participant’s ability to (i) predict upcoming stimuli and (ii) recall the shape-motor association. We found that response times decreased during the recall blocks of session one (1.07 s on run one to 1.00 s on run five; Fig. 2b, center) and of session two (from 0.97 s on run one to 0.89 s on run eight; Fig. 2b, right), demonstrating improvement in prediction and/or recall speed.

### Representational Structure in Motor Cortex Follows Response Patterns

Given the behavioral evidence for participant learning, we asked how motor response, shape, and graph were represented in the brain. In what follows, we will describe these three domains in turn. Prior work suggests that we should be able to decode consistent, subject-specific patterns of activity associated with each of the 15 one- and two-finger motor responses in both primary and somatosensory motor cortices [26]–[28]. We hypothesized that we could reliably decode the associated motor movement from the neural pattern on any given trial, and that the relationships between those neural patterns would be preserved across participants.

To measure the representation of stimulus information in the brain, we decoded stimulus identity from patterns of evoked BOLD activity using multi-voxel pattern analysis (MVPA). Each stimulus evokes a pattern of brain activation, which can be viewed as a point in high-dimensional space, where each dimension corresponds to a different voxel’s BOLD activation. For each of these activation patterns, we used a trained classifier to predict what trial was being experienced by the subject, and we assessed the veracity of that prediction using cross-validation to predict trials in a held-out run (see Methods). We examined classification accuracy in left, right, and a bilateral postcentral gyrus region of interest (ROI) by training a classifier on data from the respective laterality corresponding to seven of the session two recall runs, and by testing our prediction on the remaining run. Data consisted of LS-S *β* weights resulting from a contrast of the regressor for each trial to a nuisance regressor of all other trials. Because participants responded with their right hand, we expected greater classification accuracy in the left hemisphere than in the right, and we expected greater classification accuracy when utilizing both hemispheres than when utilizing one hemisphere.

As expected, we could predict the identity of the current trial with an accuracy above chance for all 31 participants. That is, the BOLD patterns evoked within each subject in response to finger movements were highly consistent across trials and highly distinct for each movement. Prediction accuracy was above chance for the left hemisphere postcentral gyrus ROI, and for a right hemisphere postcentral gyrus ROI and a bilateral postcentral gyrus ROI in all but one subject (Fig. 3a). Specifically, we found that left-hemisphere classification accuracy was significantly higher than right hemisphere classification accuracy (two-sided paired *t*-test; *t*_30_ = 6.55; *p <* 4 *×* 10^−7^). Interestingly, classifying held-out runs using data from both hemispheres did not provide a statistically greater accuracy than classifying using data from the left hemisphere alone (two-sided paired *t*-test; *t*_30_ = −0.64, *p* = 0.53). These results demonstrate that trial information was robustly represented in the BOLD signal in postcentral gyrus, and that we could decode this information with MVPA, particularly in the hemisphere contralateral to the motor response.

**Figure 3:**
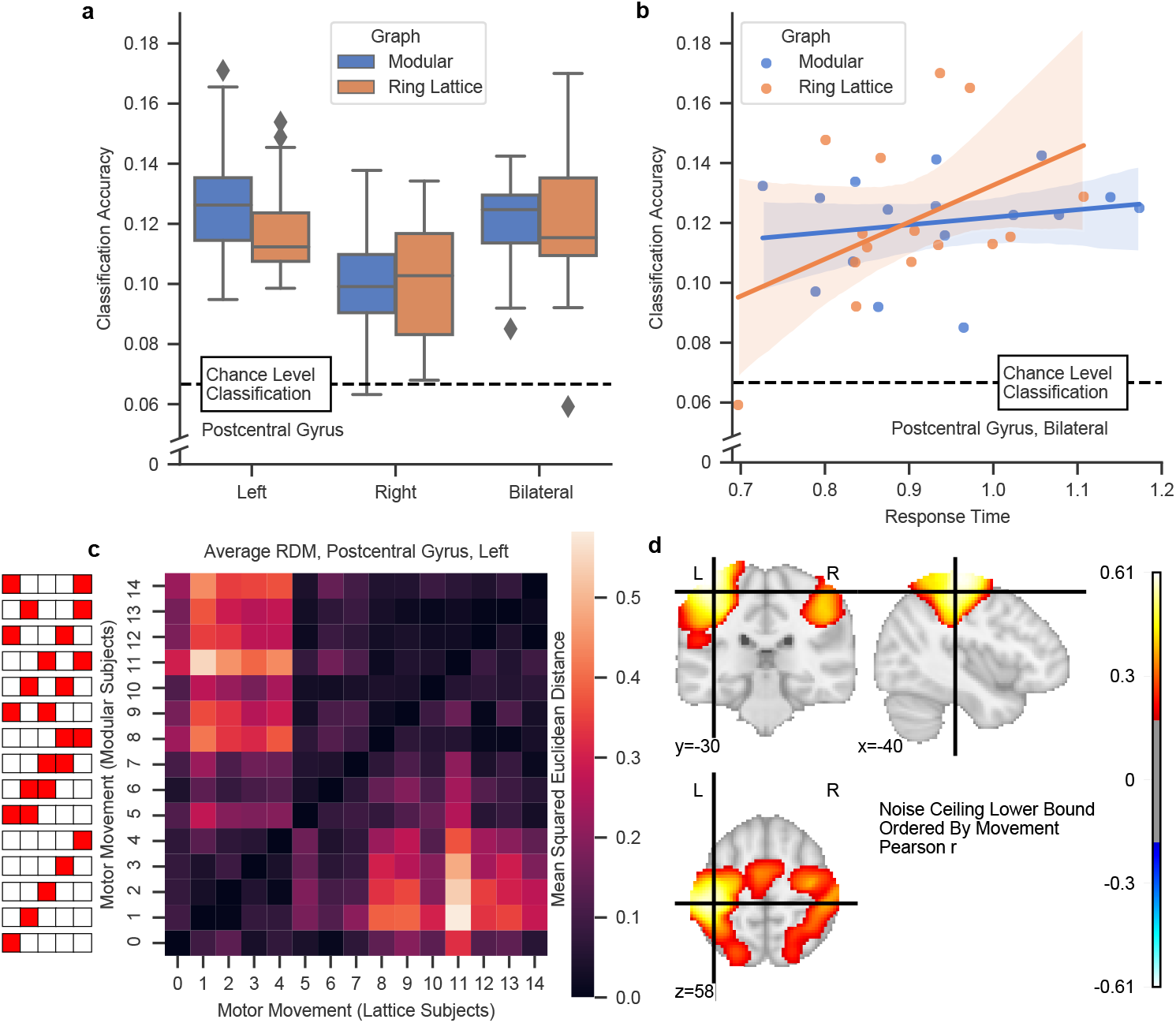
Motor responses decoded from the postcentral gyrus. (**a**) Above-chance classification accuracy for all participants in the left postcentral gyrus. Colors indicate graph condition: modular or ring lattice. Boxes show 25%, 50%, and 75% quartiles. Whiskers show the range of the data, with outliers indicated by diamond markers. The dotted line indicates chance performance of 6.67%. (**b**) SVM classification accuracy in postcentral gyrus was correlated with mean response time in the recall blocks of session two. Lines indicate linear regression fits and shaded envelopes indicate 95% confidence intervals. (**c**) Representational dissimilarity matrices (RDMs) were calculated for each subject using a cross-validated Euclidean metric, and averaged across subjects. Here we show the average RDM in left postcentral gyrus, with rows ordered by the motor command (see key at left of panel). The upper triangle presents data for the modular condition and the lower triangle presents data for the lattice condition; diagonal elements are equal to 0 for both conditions. High values indicate dissimilar patterns. (**d**) In the postcentral gyrus, the representational dissimilarity matrices were highly similar across subjects when ordered by motor response. Here we ran a searchlight to identify regions that displayed consistent representations across subjects. Shown is the average Pearson correlation coefficient, *r*, when the RDM in the neighborhood of a voxel was correlated with the average RDM of all other subjects in that neighborhood. We observed two regions of high consistency: one centered on motor cortex (shown here), and one centered on the cerebellum. The searchlight was thresholded at *r* = 0.18 to control the FWER at *p <* 0.05.

While robust, we observed that classification accuracy varied from 5.9% to 17% across subjects in the bilateral postcentral ROI. What drives these differences in classification accuracy? We hypothesized that individual differences in task performance effected differences in classification accuracy. We therefore asked whether our ability to classify trials for a participant was dependent on that participant’s mean response time or mean response accuracy. To answer this question, we used the participant behavior on the recall blocks in session two, which represented post-training task performance (Fig. 3b). We found that response time was a significant predictor of classification accuracy, such that classification accuracy was higher for participants with higher average response times than for participants with lower average response times (OLS; *β* = 0.0191, *SE* = 0.009, *t*_27_ = 2.13, *p <* 0.043). We did not observe a relationship between classification accuracy and response accuracy (OLS; *β* = −0.06, *SE* = 0.081, *t*_27_ = −0.73, *p* = 0.47). Given that the postcentral gyrus encodes information about motor movements, it is not surprising that the timing of those movements impacted our ability to decode their evoked activity.

Our classification results suggested that evoked patterns of activity were highly reliable across runs within single subjects. Prior results have suggested that activation patterns vary appreciably between subjects [27], but that the relationships *between* activation patterns are conserved. That is, the distance between representations should be consistent between subjects, even if the specific voxel-wise patterns differ. To assess the preservation of inter-representation differences across subjects, we estimated a representational dissimilarity matrix (RDM) for each subject using a cross-validated Euclidean metric (see Methods). In this matrix, we let *i, j* index pairs of movements, and we let the *ij*-th matrix element indicate the dissimilarity of movement *i*’s representation from movement *j*’s representation. We then averaged the RDMs across participants (Fig. 3c), separately for the two graph conditions (modular and ring lattice). We found that differences in motor representations were preserved across participants and graph conditions, and reflected the movements involved: representations of two-finger movements were consistently more dissimilar to representations of one-finger movements than they were to themselves (two-sided independent *t*-test; *t*_93_ = 9.45, *p <* 3 *×* 10^−15^), and likewise one-finger movement representations were more dissimilar to two-finger movements than they were to themselves (independent *t*-test; *t*_58_ = 4.70, *p <* 2 *×* 10^−5^). These data demonstrate not only that subjects exhibited reliable patterns of activity evoked by each motor response across the experiment, but also that the relationship between those patterns was reliable between subjects within postcentral gyrus.

Although we chose to consider the postcentral gyrus motivated by prior work, we also performed an exploratory analysis to identify other regions whose RDMs were highly similar across participants. For each participant, we conducted a whole-brain searchlight where for each voxel we computed the mean RDM in the neighborhood of that voxel, thereby producing a 105-dimensional vector for each voxel composed of the upper right entries in the 15-by-15 RDM. We then calculated the correlation coefficient between that vector and the mean vector of the same voxel in the remaining 30 participants. Finally, we took the mean of these correlations across all subjects. This process is similar in principle to that of estimating a lower bound on the noise ceiling [29]. The resulting map should highlight voxels that displayed similar representational dissimilarity patterns across participants, without making any assumptions about the representations themselves or about the dissimilarity pattern. We found two regions that exhibit high intersubject similarity after thresholding to control the FWER at *p <* 0.05 (see Methods): a region encompassing both pre- and postcentral gyrus bilaterally (Fig. 3d) and a region in the cerebellum (Supplemental Fig. S2). Thus by constructing an RDM organized by movement, but not by shape or node identity, we identify three regions (precentral gyrus, postcentral gyrus, and cerebellum) which exhibited reliable cross-subject relationships between evoked patterns of activity. These results suggest that by organizing the RDM by either shape or node, we we might find similar organizational patterns specific to visual or graph-level properties of trials.

### Representational Structure in Visual Cortex

Following our decoding of motor responses, we asked whether visual regions housed reliable encodings of shape identity both within and between participants. Our stimuli were chosen to produce differentiable representations in the lateral occipital cortex (LOC) [30]. In prior work, the LOC was shown to exhibit task-dependent decorrelations in the neural representations of shape stimuli [31]. We therefore hypothesized that if the architecture of the graph altered the perception or encoding of stimuli, then that dependence would manifest in LOC. Using data from a separate localizer run, we were able to localize the LOC for each subject by applying a general linear model (GLM) with the following contrast: BOLD responses to a novel set of shape stimuli versus BOLD responses to phase-scrambled versions of those shapes. Note that these novel shapes were generated using the same process as that for the shapes in the training and recall blocks. We then intersected the two largest clusters from the localizer with a surface-defined LOC ROI and ordered the voxels in this intersection by their *z*-statistics from the localizer contrast. We chose the 200 voxels with the highest *z*-scores from each hemisphere to define left and right ROIs, and we chose the top 600 voxels to define a bilateral ROI. We then studied whether stimulus coding in this ROI exhibited similar properties to the motor representations we could reliably decode in postcentral gyrus.

As before, we first wished to verify that trial-by-trial stimulus representations were consistent within subjects. We found that we could predict stimulus identity on a held-out run remarkably well across all participants and ROI choices (Fig. 4a). We averaged the classification accuracy across all 8 folds and observed that it was greater than chance (6.67%) for all participants. We also observed higher classification accuracy in participants trained on the modular graph than in participants trained on the lattice graph (Mixed ANOVA, *F* (1, 29) = 14.14, *p <* 8 *×* 10^−3^). These data demonstrate that stimulus identity could be reliably decoded within-subject from activity within LOC.

**Figure 4:**
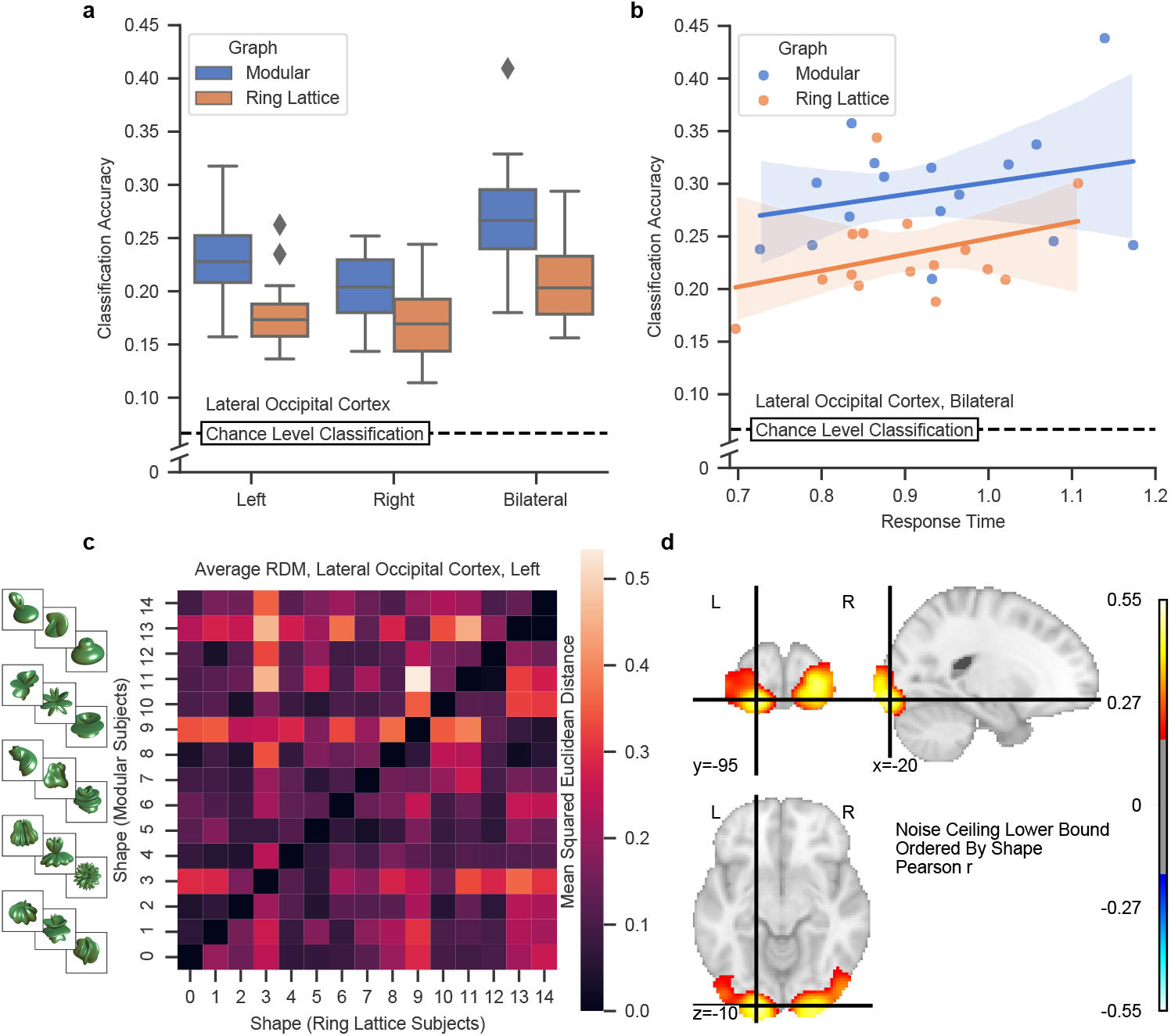
Shape stimuli decoded from the lateral occipital cortex. (**a**) Above-chance classification accuracy for all participants in left, right, and bilateral LOC (lateral occipital cortex) ROIs. Colors indicate graph condition: modular or ring lattice. Classification accuracy differs significantly between conditions (Mixed ANOVA, *F* (1, 29) = 14.14, *p <* 8 *×* 10^*−*3^). Boxes show 25%, 50%, and 75% quartiles. Whiskers show the range of the data, with outliers indicated by the diamond markers. Dotted line indicates chance performance of 6.67%. (**b**) In LOC, graph type predicted SVM classification accuracy (OLS, *t*_27_ = 2.15, *p <* 0.042, 95% CI: 0.002 to 0.104), whereas response time was not a significant predictor (OLS, *t*_27_ = 1.13, *p <* 0.27, 95% CI: -0.019 to 0.067). Lines indicate linear regression fits and shaded envelopes indicate 95% confidence intervals. (**c**) Representational dissimilarity matrices (RDMs) were calculated using a cross-validated Euclidean metric and then averaged across subjects. Here we show the average RDM in left LOC, with rows ordered by the shape stimulus (see key at left of panel). The upper triangle presents data for the modular condition and the lower triangle presents data for the lattice condition; diagonal elements are equal to 0 for both conditions. High values indicate dissimilar patterns. (**d**) In the visual cortex, the RDMs were highly similar across subjects when ordered by stimulus shape. Here we ran a searchlight to identify regions that displayed consistent representations across subjects. Shown is the average Pearson correlation coefficient, *r*, when the RDM in the neighborhood of a voxel was correlated with the average RDM of all other subjects in that neighborhood. We observed one region of high consistency, which was centered bilaterally on LOC and extended posteriorly to early visual cortex. The searchlight was thresholded at *r* = 0.18 to control the FWER at *p <* 0.05.

To evaluate whether classification accuracy differed by graph, we fit a linear regression model that predicted classification accuracy from both response time and graph condition (Fig. 4b). We found that graph condition (modular *versus* ring lattice) was a statistically significant predictor of classification accuracy (OLS, estimated increase of 5.3% for the modular graph, *t*_27_ = 2.15, *p <* 0.042, 95% CI: 0.002 to 0.104). This result suggests that there are fundamental differences in learned representations between the two training regimens. In contrast to our findings in the postcentral gyrus, the classification accuracy in LOC was not predicted by response time (OLS, *β* = 0.024, *t*_27_ = 1.13, *p <* 0.27, 95% CI: -0.019 to 0.067). Thus we find that graph structure induced a significant difference in our classification accuracy of stimuli, and that difference, as is evident in Fig. 4b, was not driven by response time differences.

Given the reliable patterns of activity in LOC, we next computed a mean RDM in the LOC across subjects as in Fig. 3c, but arranged the RDM by shape rather than by motor command. Crucially, this RDM was not a reordering of the mean motor response RDM, as the correspondence between shapes and motor responses was distinct for each subject. In computing the mean RDM, we averaged data separately for modular and ring lattice conditions. Across both graphs, we found a consistent distance relationship between shape representations (note the symmetry between the upper and lower triangles of Fig. 4c). This finding suggests that the sensitivity of the LOC to individual shape features was shared across subjects.

The analyses we have described thus far focused on the LOC. Next, similar to Fig. 3d), we performed an exploratory analysis to determine whether other areas of the brain displayed RDMs that were consistent across subjects when the RDMs were arranged by shape. We computed the correlation between a single subject’s shape RDM and the average shape RDM across all other subjects, in the neighborhood of each voxel (Fig. 4d). As before, we took the mean of this correlation estimated for all subjects, and applied a threshold to control the FWER at *p <* 0.05 (see Methods). We observed a single region of high between-subject similarity. As expected, this region was centered on the LOC and extended posteriorly through early visual cortex. Interestingly, we observed no other regions where RDMs were consistent between subjects when arranged by shape. These results largely mirror our findings with motor response activity patterns. However, here we demonstrate that by simply rearranging our RDM we can instead isolate relationships between activity driven by visual properties of the stimuli. Intriguingly, we observe a difference in decoding accuracy between graph conditions, suggesting that graph structure modulates the encoding of individual stimuli in LOC.

### Graph Effects

By organizing the RDMs according to stimulus shape and motor command, we identified two sets of regions in which patterns of representational dissimilarity were shared across participants. This inter-subject similarity was likely driven by aspects of stimulus identity: one- and two-finger movements for the motor response, and aspect ratio or curvature for stimulus shapes. In a final analysis, we turned to the last dimension along which representations could be ordered in our experiment: that of the graph. In particular, we reorganized the RDM by graph node and observed no clear region in which representational structure was shared across participants (Fig. 5a). This variation is perhaps intuitive, as graph structure is experienced indirectly through a sequence of steps between stimuli, and also variably in the sense that each participant experienced a different walk through the network. Hence, it is natural to expect high variability in how each participant encodes the graph structure.

**Figure 5:**
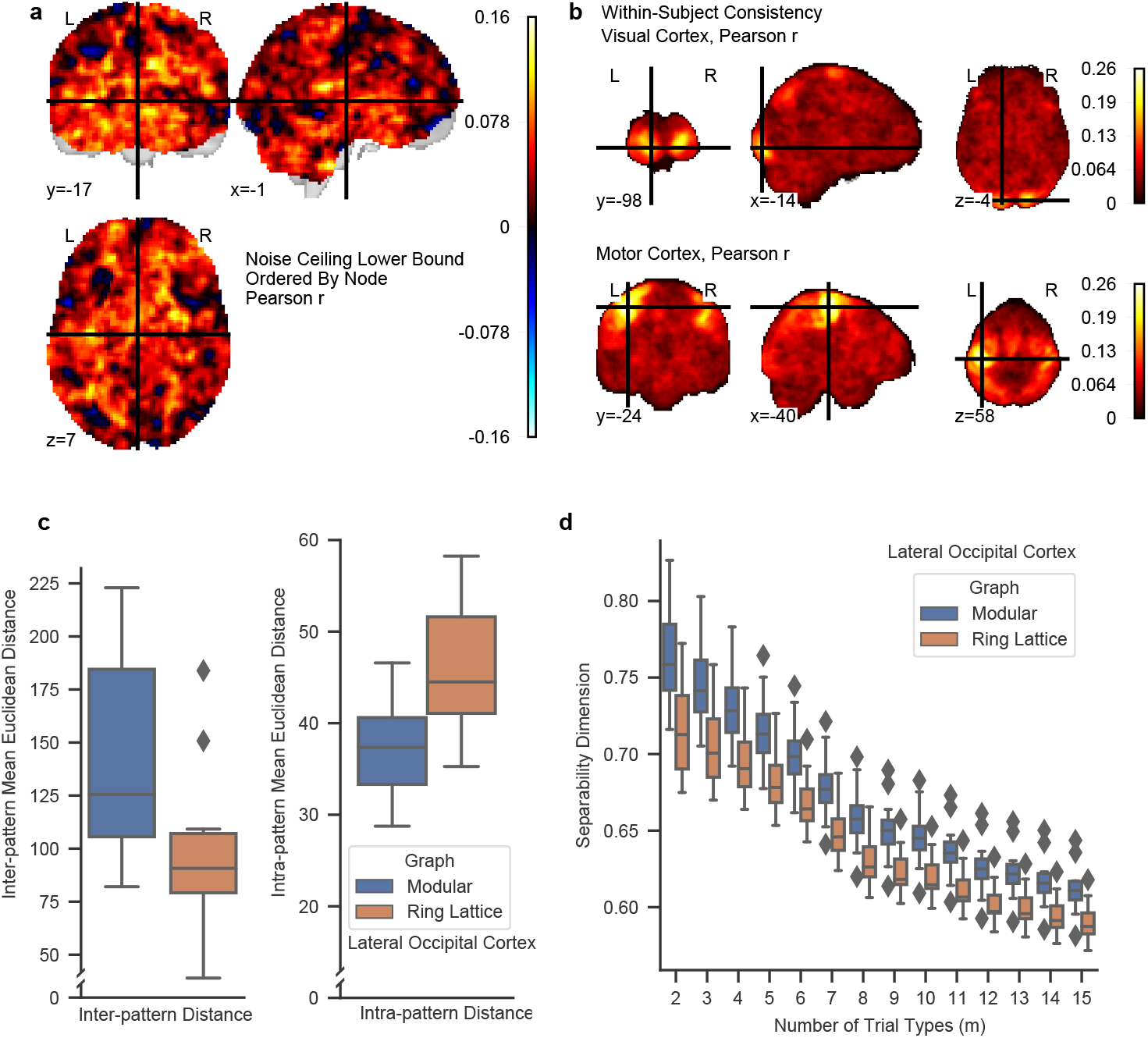
Effects of graph structure on representation. (**a**) We did not observe similar representations of graph structure across participants. Here we ran a searchlight to identify regions that showed high similarity in how graph nodes were represented relative to one another. Shown are the mean values when the RDM in the neighborhood of a voxel was correlated with the average RDM of all other subjects in that neighborhood. We computed this quantity separately for both graph conditions and then averaged the two maps. (**b**) Two regions exhibited heightened within-subject consistency of representational dissimilarity matrices: visual cortex (*top*) and motor cortex (*bottom*). Both panels present the mean Pearson’s correlation coefficient, *r*, between RDMs among runs for each subject. (**c**) Subjects in the modular graph condition showed higher average distance between representations of different trial types (*left*) and lower average distance between representations of the same trial type (*right*) than subjects in the lattice condition. The inter-representation distance was the mean of the off-diagonal elements of the Euclidean distance RDM for each subject. Higher values indicate that trial type representations were more distinct from one another. Meanwhile, the intra-representation distance measures how invariant a representation is across repetitions of the same trial type. Low values indicate consistent evoked activity. (**d**) Here we show the separability dimension of the representations in LOC. We observed a consistently higher separability dimension in the modular condition than in the ring lattice condition.

High intersubject variability can exist alongside high intrasubject reliability. To understand the potential relation between the two, we measured intrasubject reliability by the consistency of RDMs across runs within each subject. We performed a similar procedure to our earlier searchlight, with the one key difference being that we estimated the correlation between the RDM for each run and the mean of the RDMs for the other runs, in the neighborhood of each voxel. This approach provided us with a consistency map for each subject, and similar to before, we averaged these consistency maps across subjects (Fig. 5b). We observed two regions of high consistency that overlap with the motor and visual regions previously observed to exhibit high inter-subject reliability: that is, all regions where we observed high intra-subject reliability were also the regions in which we observed high inter-subject reliability. This finding suggests that any consistent representation of graph structure may spatially overlap with stimulus shape or motor response encodings.

Consistent with this suggestion, we observed that graph type was a significant predictor of classification accuracy in LOC (Fig. 4a,b), whereby classification accuracy was greater in the modular condition than in the lattice condition. One possible explanation for this observation is that trial types were more distinguishable from one another due to more distinct representations: when a subject experienced the modular graph, the representation of shape one may simply have been further away from the representation of shape two in representational space. To evaluate the validity of this possible explanation, we calculated the average distance between representations in the bilateral LOC ROI by computing a Euclidean distance RDM for each run, and then taking the mean of the upper-right off-diagonal elements for each subject (Fig. 5c, left). We observed that discriminability was significantly greater in the modular graph than in the lattice graph (two-sided independent *t*-test; *t*_29_=3.04, *p*=0.005). That is, the modular condition induced more separable mean activity patterns between pairs of stimuli than did the lattice condition.

Another possible explanation for the greater classification accuracy observed in the modular graph is that trial-to-trial replicability may be higher when a subject experiences the modular graph: a given shape may evoke a more consistent neural representation on each appearance of that shape. To evaluate this possibility, we calculated the average consistency of representations in the same bilateral LOC ROI by taking the mean representation for each run and trial type, and then calculating the mean distance of that representation from the representations of the other runs (Fig. 5c, right). We observed that patterns were more consistent in the modular graph than in the lattice graph, as evident in lower distances between trial repetitions (two-sided independent *t*-test; *t*_29_=3.60, *p* < 0.002). Taken together, these two findings suggest that the modular graph leads to more distinguishable and consistent trial representations, which in turn could drive the heightened classification performance.

The observed increase in neural discriminability in LOC suggests that behavioral discriminability might similarly differ between graph conditions. To test for this possibility, we recruited an additional group of subjects and trained them on the same set of stimuli associated to either a modular or lattice graph. We then tested whether subjects could more quickly discriminate between stimuli presented on the modular graphs than between the same stimuli presented on the lattice graph (Supplemental Fig. S3). We found that response times when discriminating between shape pairs were highly correlated with their representational distance in LOC (Mixed Effects Model, *t*_93_=4.33, *p* < 0.0001). Notably, though, we did not observe a significant difference based on graph type (Mixed Effects Model, mean=0.07 ms, 0.076280, *t*_48_=0.866, *p* < 0.392). These behavioral results suggest that while subjects in the modular condition exhibited more distinguishable LOC activity patterns than did subjects in the lattice condition, these differences in stimulus representation do not directly relate to behavioral distinguishablity of stimuli.

A third possible explanation for the greater classification accuracy observed in LOC in the modular graph is the dimensionality of the representation. If one graph topology allows a participant to encode representations in a higher dimensional space, then those representations could also be made more distinct from one another. Note that our ability to classify trials using an SVM relies on finding a hyperplane separating the trial types. Intuitively, if the trial types lie in a higher-dimensional space, then finding a separating hyperplane becomes an easier task than if the trial types lie in a low-dimensional space. To determine whether dimensionality plays a role in LOC’s representations, we computed a quantity called the *separability dimension*, which measures the fraction of binary classifications that can be implemented using our trial types (see Refs. Rigotti, Barak, Warden, *et al*. [32] and Tang, Mattar, Giusti, *et al*. [33] and Methods for details). Consistent with our intuition, we found that dimensionality was higher in the modular graph condition than in the ring lattice condition (Fig. 5d). Interestingly, this difference was apparent in LOC where we observed a difference in classification accuracy between graph conditions, but not present in the postcentral gyrus where we observed no difference in classification accuracy between graph conditions (Supplemental Fig. S1).

## Discussion

Humans deftly learn the networks of interactions underlying complex phenomena such as language or music [2]. Behavioral and neural evidence demonstrates that the way we learn and represent stimuli is modulated by the temporal structure in which we experience those stimuli [34], [35]. Moreover, recent studies indicate that humans may learn to represent the interaction network itself. For example, when stimuli are grouped into clusters, a phenomena commonly observed in natural settings [9], learner response times to those stimuli are modulated by whether adjacent stimuli are situated in the same cluster, even when all transition probabilities (and thus predictive information) are held constant [23]. This effect is shown to be robust to types of stimuli [36] and significant topological variation [37]. Moreover, learners respond more swiftly to a sequence drawn from a topology with such clusters compared to a topology without such clusters [7]. Collectively, these findings suggest that humans can learn properties of graph structure such as modular organization, and can leverage that structure to create more effective predictions. Converging neuroimaging evidence suggests that neural representations of stimuli encode properties of the interaction networks, including cluster identity [24] and graph distance between items [25], [38]. To date, however, it remains unclear how such graph-induced representation structure differs when the same elements are arranged in a distinct organizational pattern.

In the current study, we addressed this question by training human subjects to respond to sequences of paired visual and motor stimuli. We then tested whether representations of those stimuli differed when we performed a controlled manipulation of the sequence. Specifically, each subject learned a sequence based on either a modular graph or a ring lattice graph, in both cases with no explicit knowledge of the underlying graph. Subjects underwent one day of training followed by a second day in which they were asked to produce the motor response associated with each visual stimulus in the sequence. We found that we could reliably decode stimulus identity from the neural representation in both postcentral gyrus and lateral occipital cortex. Further, we observed consistent representation structure of our stimulus set across subjects in these regions, when comparing motor response in postcentral gyrus and stimulus shape in lateral occipital cortex.

Within lateral occipital cortex, but not postcentral gyrus, our classification ability was modulated by graph type, with the modular graph leading to significantly higher classification scores than the ring lattice graph. This classification difference due to graph was accompanied by changes in the representation structure: modular representations were both more consistent across trials and more separable between different trial types than were representations from the ring lattice graph. Finally, we found that representations in lateral occipital cortex from the modular graph exhibited higher separability dimension than did those from the ring lattice graph. Taken together, these results show how graph structure can modulate learned neural representations of stimuli, and that modular structure induces more effective (discriminable) representations of stimuli than does ring lattice structure.

### Modular graphs enable effective neural representations

There is strong theoretical justification to expect more robust representations to arise from the modular graph than from other non-modular graphs. Using information theory, prior work has shown that graph organizations characterized by hierarchical modularity facilitate efficient transmission of information [9]. Perhaps due to that functional affordance, this type of organization is commonly observed in many real-world networks such as those of word transitions, semantic dependencies, or social relationships. The informational utility of hierarchically modular networks arises in part from the fact that such networks exhibit high entropy, thereby communicating large amounts of information. It also arises in part from the expectations that humans form about clustered structures: preferentially predicting transitions that remain within a cluster, thereby accurately foreshadowing upcoming stimuli [8]. Here in our study, the modular graph composed of three clusters of five nodes is more highly clustered than the ring lattice graph. This fact can be clearly seen visually, but it can also be quantified by the clustering coefficient, which is defined as the fraction of a node’s neighbors that are connected to one another. The average clustering coefficient of the modular graph is 0.7 and of the ring lattice graph is 0.5. Indeed, we found that participants formed more robust representations of stimuli from the modular graph than from the lattice graph. In other words, the modular graph is more similar to graphs encountered in the real world, to which our expectations are tuned, and is also better capable of conveying its structure.

### Neural representations from modular graphs are high dimensional

We observed that the neural representations developed during exposure to a sequence drawn from a modular graph tend to be high dimensional. Prior work has shown that high dimensionality of neural representations is associated with faster learning and more efficient stimulus embedding[33]. Here, we extend the current knowledge in the field by demonstrating that more efficient *graph structures* lead to higher dimensional neural representations. It is possible that this association between topology and dimensionality arises from the fact that the modular graph can be neatly separated into clusters; coding by cluster could allow the brain to use a high-dimensional embedding. The nature of this higher-dimensional encoding remains a topic for future study, and could be pursued by investigating the learning of a variety of graphs with and without hierarchical structure.

### Graph-dependent adaptation decorrelates shapes

Our observation—that graph topology can be used to decorrelate neural responses to shape—is of particular relevance to the study of neural adaptation. An oft-hailed mechanism of neural adaptation is the decorrelation of neural responses among stimuli [39], [40]. Such decorrelation is thought to allow the brain to become attuned to task-relevant distinctions between stimuli [41]. Prior work has observed decorrelation in BOLD responses to visual stimuli in the lateral occipital cortex Mattar, Olkkonen, Epstein, *et al*. [31]. Specifically, this study found that training on two classes of stimuli caused representations evoked by the two classes in the lateral occipital cortex to decorrelate from one another. Such increased neural separability of stimuli may support behavioral discrimination between important stimulus classes.

Here in our study, all subjects learned the same set of 15 previously unseen shapes. Basic visual properties of the shapes are expected to drive the BOLD response in the lateral occipital cortex [30]. However, if these properties alone drive the BOLD response, then we would expect similar representational structure between activity patterns in both modular and ring lattice subjects. Instead, we found that we could more accurately classify stimuli from neural representations when participants experienced sequences determined by the modular graph than when they experienced sequences determined by the ring lattice, suggesting that BOLD responses are driven by non-visual factors. This classification difference was accompanied by two observed differences in pattern distance: First, activation for a given trial type was more stable across trials in the modular graph than in the ring lattice graph, as evidenced by the lower mean pattern distance between trial repetitions. Second, activation was more distinct between trials of different types: the average distance between different trial types was higher in the modular graph than in the ring lattice graph. In summary, we observe that graph structure drives a difference in representation fidelity, inducing both more reliable and more separable patterns.

### Motor cortex displays strong organizational patterns

The approach that we use here—representational similarity analysis or RSA—has been commonly used to understand how neural representations of stimuli are encoded. Rather than asking which neurons or voxels encode a stimulus, we can study how the patterns of neurons or voxels relate across stimuli. Further, we can compare neural distances to those predicted by real-world stimulus properties, elucidating which real-world properties drive the brain’s organization. For instance, the same single-digit finger movement in different people elicits vastly different patterns of neural activation in primary motor cortex, while the relationship between activity patterns of different fingers is highly conserved across people [27]. Prior work has demonstrated that everyday usage patterns predict motor representations better than musculature, a result that fundamentally underscores the importance of *relationships* rather than the movements themselves. Similar findings have also been reported for motor sequences[28].

Here in our study, we recorded neural activity of both single- and multi-finger movements, and found that these relationships were highly preserved between participants and between graphs (Fig. 3c). This observation raises an intriguing question for further study: Why is the difference in representational dimensionality between conditions observed in LOC but not in motor cortex? Strikingly, in the postcentral gyrus we found a clear separation of one-finger and two-finger response patterns, where one-finger response patterns were similar to other one-finger response patterns, and two-finger response patterns were similar to other two-finger response patterns. We note that our ROI encompasses a larger area than that used in prior work [27], and studies of multi-finger sequences, while distinct from multi-finger movements, have observed that these increasingly complex movements elicit distinct representations in areas further from the central sulcus [28]. Further investigation is necessary to determine whether (and to what degree) the dissimilarity patterns we observed are influenced by differences in cortical location of one-finger and two-finger motions.

## Limitations

Prior work on learning modular structure has found that graph information such as cluster identity is represented in the medial temporal lobe (MTL) [4], [24], an area also shown to encode distances between items [25]. In the current study, we did not observe any graph-dependent effects in the MTL, but instead in the lateral occipital cortex. A number of factors could give rise to this difference. First, prior studies, to our knowledge, did not show evidence that graph structure was directly encoded. Metrics such as distance and cluster identity pool over many trials, and may allow for greater power to detect graph-related effects in the presence of noise. Indeed, the MTL is prone to low signal-to-noise-ratio fMRI signals due to anatomy [42]. This can be improved by imaging with a reduced field of view, but at a limitation of our ability to infer graph effects elsewhere in the brain. Additionally, our task is relatively complex. Behavioral work on graph learning has frequently required participants to learn image orientations [3], [23] or to anticipate motor responses [7] but here participants must learn a mapping between both images and motor responses. Indeed, participants are still learning the shape-motor response pairings up through the end of the second session, as response accuracy continues to increase (Fig. 2b, *right*). The complexity of the task, requiring participants to learn not only visual stimuli but the stimulus-motor response pairings as well, could require additional training for the MTL to detect temporal regularities. Given that plasticity in the lateral occipital cortex can be driven by demands of stimulus discriminability [31], additional training may be needed for the hippocampus to extract temporal statistics.

## Conclusion

In this study, we expand upon prior behavioral evidence in graph learning, where graphs with modular structures produce faster response times in learners, suggesting that those learners can form more accurate predictions. Learners were successfully able to learn and recall a mapping of 15 novel shapes to motor movements, combining behavioral paradigms from prior graph learning work. We observed that modular graphs enable more consistent and distinct representations when compared to a ring lattice, and in turn allow better prediction of what a participant is responding to using machine learning. This representational difference was present in the lateral occipital cortex, a region sensitive to properties of the visual stimuli used in this task, and was accompanied by increased dimensionality of representations, a feature known to accompany effective learning. Our results motivate future work to better understand the nature of these representational changes, as well as generalizability to broader sets of graph structures.

## Citation Diversity Statement

Recent work in several fields of science has identified a bias in citation practices such that papers from women and other minority scholars are under-cited relative to the number of such papers in the field [43]–[51]. Here we sought to proactively consider choosing references that reflect the diversity of the field in thought, form of contribution, gender, race, ethnicity, and other factors. First, we obtained the predicted gender of the first and last author of each reference by using databases that store the probability of a first name being carried by a woman [47], [52]. By this measure (and excluding self-citations to the first and last authors of our current paper), our references contain 13.18% woman(first)/woman(last), 7.12% man/woman, 26.79% woman/man, and 52.92% man/man. This method is limited in that a) names, pronouns, and social media profiles used to construct the databases may not, in every case, be indicative of gender identity and b) it cannot account for intersex, non-binary, or transgender people. Second, we obtained predicted racial/ethnic category of the first and last author of each reference by databases that store the probability of a first and last name being carried by an author of color [53], [54]. By this measure (and excluding self-citations), our references contain 6.03% author of color (first)/author of color(last), 14.19% white author/author of color, 22.81% author of color/white author, and 56.98% white author/white author. This method is limited in that a) names and Florida Voter Data to make the predictions may not be indicative of racial/ethnic identity, and b) it cannot account for Indigenous and mixed-race authors, or those who may face differential biases due to the ambiguous racialization or ethnicization of their names. We look forward to future work that could help us to better understand how to support equitable practices in science.

## Acknowledgments

This research was primarily supported by the National Institutes of Mental Health award number 1-R21-MH-124121-01. D.S.B. would also like to acknowledge additional support from the John D. and Catherine T. MacArthur Foundation, the Alfred P. Sloan Foundation, the Institute for Scientific Interchange Foundation, and the Army Research Office (Grafton-W911NF-16-1-0474 and DCIST-W911NF-17-2-0181). The content is solely the responsibility of the authors and does not necessarily represent the official views of any of the funding agencies.

## Author Contributions

A.E.K., G.K.A., and D.S.B. conceived the project and planned the experiments and analyses. A.E.K., K.S., E.B.H., and N.N. performed the experiments and analyses. S.L. replicated the analyses and results. A.E.K. and D.S.B. wrote the manuscript and Supplementary Information. K.S., E.B.H., N.N., G.K.A., and D.S.B. edited the manuscript and Supplementary Information.

## Methods

### Task

#### Stimuli

Visual stimuli were displayed on either a laptop screen or on a screen inside the MRI machine using PsychoPy v3.0.3 and consisted of fifteen unique abstract shapes generated by perturbing a sphere with sinusoids using the `MATLAB` package `ShapeToolbox` (Fig. 1a, *left*). Each shape consisted of two sinusoidal oscillations. Each oscillation could vary in amplitude (either 0.2 or 1), angle (0, 30, 60, or 90 degrees), frequency (2, 4, 8, 10, or 12 cycles/2*π*), and the second oscillation could also vary in phase relative to the first oscillation (0 or 45 degrees). Out of the set of 3200 permutations, we selected a set of 15 visually distinct shapes for the main experiment. A second set of 15 visually distinct shapes was selected from the same set for the localizer. The same set of stimuli was used for all participants, albeit in different order. Five variations of each shape were also created through scaling and rotating (scale/rotation pairs were 100%/0 degrees, 95%/5 degrees, 100%/10 degrees, 95%/15 degrees, and 100%/20 degrees) and the appearance of variations was balanced within task runs so as to capture neural activity invariant to these properties.

Each shape was paired with a unique motor response. Possible motor responses spanned the set of all fifteen one- or two-button chords on a five-button response pad (Fig. 1a, *right*). The five buttons on the response pad corresponded to the five fingers on the right hand: the leftmost button corresponded to the thumb, the second button from the left corresponded to the index finger, and so on. In addition to the shape cue, motor responses were explicitly cued by a row of five square outlines on the screen, each of which corresponded to a button on the response pad. Squares corresponding to the cued buttons turned red; for instance, if the first and fourth squares turned red, a correct response consisted in the thumb and ring finger buttons being pressed.

Participants were assigned one of two graph types: modular or ring lattice. Both graph types were composed of fifteen nodes and thirty undirected edges such that each node had exactly four neighbors (Fig. 1b, *left*). Nodes in the modular graph were organized into three densely connected five-node clusters, and a single edge connected each pair of clusters. Each node in a participant’s assigned graph was associated with a stimulus-response pair (Fig. 1b, *right*). While the stimulus and response sets were the same for all participants, the stimulus-response-node mappings were random and unique to each participant. This mapping was consistent across runs for an individual participant. The order in which stimuli were presented during a single task run was dictated by a random walk on a participant’s assigned graph; that is, an edge in the graph represented a valid transition between stimuli. No other meanings were ascribed to edges. Walks were randomized across participants as well as across runs for a single participant. Two constraints were instituted on the random walks: (1) all walks were required to visit each node at least ten times, and (2) walks on the modular graph were required to include at least twenty cross-cluster transitions.

#### Pre-Training

Prior to entering the scanner, participants performed a “pre-training” session during which stimuli were displayed on a laptop screen and motor responses were logged on a keyboard using the keys ‘space’, ‘j’, ‘k’, ‘l’, and ‘;’. Participants were provided with instructions for the task, after which they completed a brief quiz to verify comprehension. To facilitate the pre-training process, shape stimuli were divided into five groups of three. First, participants were simultaneously shown a stimulus and its associated motor response cue, and then were asked to press the associated button(s) on the keyboard. Next, participants were shown the stimulus alone and asked to perform the motor response from memory. If they did not respond within two seconds, the trial was repeated. If they responded incorrectly, the motor response cue was displayed. Once the participant responded correctly, the same procedure was repeated for the other two stimuli in the subset. Then, participants were instructed to respond to a sequence of the same three stimuli, presented without response cues. The sequence was repeated until the participant responded correctly for 12 consecutive trials. This entire process was repeated for each three-stimulus subset. The three stimuli in a given subset were always chosen so that none were adjacent to one another in the graph dictating the order of the sequence used during the “Training” stage (see below).

#### Training

During the first scanning session, participants completed five runs each consisting of a “training” and “recall” block inside the MRI scanner. A training block consisted of 300 trials and was designed to elicit learning of graph structure as in Ref. [7]. During each trial, participants were presented with a visual stimulus consisting of a row of five square outlines, each of which contained the same abstract shape (Fig. 1c). Five hundred milliseconds later, one or two of the squares turned red. Using a five-button controller, participants were required to press the button(s) corresponding to the red squares on the screen. If a participant pressed the wrong button(s), the word “Incorrect” appeared on the screen. The following stimulus did not appear until the correct buttons were pressed. Participants were instructed to respond as quickly as possible. Sequences were generated via random walks on the participants’ assigned graphs.

#### Recall (Session One)

At the end of each training run, participants completed 15 additional “recall” trials during which each shape was presented without a motor response cue, requiring participants to recall the correct response. This block was intended to measure stimulus-response association learning, as well as to prepare participants for the extended recall phase in session two. A brief instructional reminder was displayed before the start of the block. On each trial, the target stimulus was displayed in a single large square in the center of the screen. Participants were given two seconds to respond to each stimulus; if they did not respond within this time, the trial was marked as incorrect. If they responded incorrectly, the trial was marked as incorrect and the motor response cue was displayed below the shape for the remainder of the two seconds. If they responded correctly, the outline of the square containing the shape turned green. The orderings of all recall trials were also determined by walks on the participants’ assigned graphs. Trials were shown with a variable ITI between presentations. The ITIs in each block varied between two and four seconds (mean=2.8 s, total 42 s) with an additional six second delay separating the end of the last recall trial in a block from the start of the first training trial in the subsequent run.

#### Recall (Session Two)

During the second session, participants completed eight recall runs, each comprised of four recall blocks of 15 trials, for 60 trials per run. These extended recall runs were designed to allow measurement of the neural patterns of activation associated with each of the stimuli. Each constituent recall block was identical to the recall blocks presented in session one. To generate a continuous walk for each run (across the four blocks), the first block in each run began on a randomly selected node, and each subsequent block in the run was rotated to be contiguous with where the last set ended. This process ensured that the full 60 trials within a block constituted a valid random walk, despite not being generated as a single walk. Blocks 1, 2, and 4 followed a random walk, whereas the third block followed a Hamiltonian walk, in which every node was visited exactly once. Across the full run of 60 trials, each node was required to be visited at least twice. If not, the walk for the run was discarded and a new walk was generated. For the modular graph, the Hamiltonian walk proceeded clockwise on runs 1, 3, 5 and 7 and counter-clockwise on runs 2, 4, 6, and 8. On the ring lattice, the Hamiltonian walk was not generated with any directionality.

### Imaging

#### Acquisition

Each participant underwent two sessions of scanning, acquired with a 3T Siemens Magnetom Prisma scanner using a 32-channel head coil. In session one, two pre-task scans were acquired as participants rested inside the scanner, followed by five task scans and two post-task resting-state scans. In session two, two pre-task scans were acquired, followed by eight task scans, two post-task resting-state scans, and lastly an LOC localizer. Imaging parameters were based on the ABCD protocol [55]. Each scan employed a 60-slice gradient-echo echo-planar imaging sequence with 2.4 mm isotropic voxels, an iPAT slice acceleration factor of 6, and an anterior-to-posterior phase encoding direction; the repetition time (TR) was 800 ms and the echo time (TE) was 30 ms. A blip-up/blip-down fieldmap was acquired at the start of each session. A T1w reference image was acquired during session one using a `MEMPRAGE` sequence [56] with 1.0 mm isotropic voxels, anterior-to-posterior encoding, and a GRAPPA acceleration factor of 3 [57]; the repetition time (TR) was 2530 ms and the echo times (TEs) were 1.69 ms, 3.55 ms, 5.41 ms, and 7.27 ms. A T2w reference image was acquired during session two using the variable flip angle turbo spin-echo sequence (Siemens SPACE; [58]) with 1.0 mm isotropic voxels, anterior-to-posterior phase encoding, and a GRAPPA acceleration factor of 2; the repetition time (TR) was 3200 ms and the echo time (TE) was 565 ms. A diffusion-weighted scan was also acquired at the end of session two. The acquisition followed the ABCD protocol with 1.7 mm isotropic voxels, an iPAT slice acceleration factor of 3, and an anterior-to-posterior phase encoding direction; the repetition time (TR) was 4200 ms and the echo time (TE) was 89 ms, and b-values were 500 (6 directions), 1000 (15 directions), 2000 (15 directions), and 3000 (60 directions). The total acquisition time per scan was 7:29 min and consisted of 81 slices.

#### Preprocessing

Results included in this manuscript come from preprocessing performed using *fMRIPrep* 20.1.0 (Esteban, Markiewicz, Blair, *et al*. [59]; Esteban, Blair, Markiewicz, *et al*. [60]; RRID:SCR_016216), which is based on *Nipype* 1.4.2 (Gorgolewski, Burns, Madison, *et al*. [61]; Gorgolewski, Esteban, Markiewicz, *et al*. [62]; RRID:SCR_002502).

##### Anatomical data preprocessing

A total of 1 T1-weighted (T1w) images were found within the input BIDS dataset. The T1-weighted (T1w) image was corrected for intensity non-uniformity (INU) with `N4BiasFieldCorrection` [63], distributed with ANTs 2.2.0 [64, RRID:SCR_004757], and used as T1w-reference throughout the workflow. The T1w-reference was then skull-stripped with a *Nipype* implementation of the `antsBrainExtraction.sh` workflow (from ANTs), using OASIS30ANTs as target template. Brain tissue segmentation of cerebrospinal fluid (CSF), white-matter (WM), and gray-matter (GM) was performed on the brain-extracted T1w using `fast` [FSL 5.0.9, RRID:SCR_002823, 65]. Brain surfaces were reconstructed using `recon-all` [FreeSurfer 6.0.1, RRID:SCR_001847, 66], and the brain mask estimated previously was refined with a custom variation of the method to reconcile ANTs-derived and FreeSurfer-derived segmentations of the cortical gray-matter of Mindboggle [RRID:SCR_002438, 67]. Volume-based spatial normalization to two standard spaces (MNI152NLin2009cAsym, MNI152NLin6Asym) was performed through nonlinear registration with `antsRegistration` (ANTs 2.2.0), using brain-extracted versions of both the T1w reference and the T1w template. The following templates were selected for spatial normalization: *ICBM 152 Nonlinear Asymmetrical template version 2009c* [Fonov, Evans, McKinstry, *et al*. [68], RRID:SCR_008796; TemplateFlow ID: MNI152NLin2009cAsym], *FSL’s MNI ICBM 152 nonlinear 6th Generation Asymmetric Average Brain Stereotaxic Registration Model* [Evans, Janke, Collins, *et al*. [69], RRID:SCR_002823; TemplateFlow ID: MNI152NLin6Asym].

##### Functional data preprocessing

For each of the 18 BOLD runs per participant (across all tasks and sessions), the following preprocessing was performed. First, a reference volume and its skull-stripped version were generated using a custom methodology of *fMRIPrep*. Head-motion parameters with respect to the BOLD reference (transformation matrices, and six corresponding rotation and translation parameters) are estimated before any spatiotemporal filtering using `mcflirt` [FSL 5.0.9, 70]. BOLD runs were slice-time corrected using `3dTshift` from AFNI 20160207 [71, RRID:SCR_005927]. A B0-nonuniformity map (or *fieldmap*) was estimated based on two (or more) echo-planar imaging (EPI) references with opposing phase-encoding directions, with `3dQwarp` Cox and Hyde [71] (AFNI 20160207). Based on the estimated susceptibility distortion, a corrected EPI (echo-planar imaging) reference was calculated for a more accurate co-registration with the anatomical reference.

The BOLD reference was then co-registered to the T1w reference using `bbregister` (FreeSurfer) which implements boundary-based registration [72]. Co-registration was configured with nine degrees of freedom to account for distortions that remained in the BOLD reference. The BOLD time-series (including those to which slice-timing correction had been applied) were resampled onto their original, native space by applying a single, composite transform to correct for head-motion and susceptibility distortions. These resampled BOLD time-series will be referred to as *preprocessed BOLD in original space*, or just *preprocessed BOLD*. The BOLD time-series were resampled into several standard spaces, correspondingly generating the following *spatially-normalized, preprocessed BOLD runs*: MNI152NLin2009cAsym and MNI152NLin6Asym.

Several confounding time-series were calculated based on the *preprocessed BOLD*: framewise displacement (FD), DVARS, and three region-wise global signals. FD was computed using two formulations: absolute sum of relative motions Power, Mitra, Laumann, *et al*. [73] and relative root mean square displacement between affines Jenkinson, Bannister, Brady, *et al*. [70]. FD and DVARS are calculated for each functional run, both using their implementations in *Nipype* [following the definitions by 73]. The three global signals are extracted within the CSF, the WM, and the whole-brain masks. Additionally, a set of physiological regressors were extracted to allow for component-based noise correction [*CompCor*, 74]. Principal components are estimated after high-pass filtering the *preprocessed BOLD* time-series (using a discrete cosine filter with 128s cut-off) for the two *CompCor* variants: temporal (tCompCor) and anatomical (aCompCor). tCompCor components are then calculated from the top 5% variable voxels within a mask covering the subcortical regions. This subcortical mask is obtained by heavily eroding the brain mask, which ensures it does not include cortical GM regions. For aCompCor, components are calculated within the intersection of the aforementioned mask and the union of CSF and WM masks calculated in T1w space, after their projection to the native space of each functional run (using the inverse BOLD-to-T1w transformation). Components are also calculated separately within the WM and CSF masks. For each CompCor decomposition, the *k* components with the largest singular values are retained, such that the retained components’ time series are sufficient to explain 50 percent of variance across the nuisance mask (CSF, WM, combined, or temporal). The remaining components are dropped from consideration.

The head-motion estimates calculated in the correction step were also placed within the corresponding confounds file. The confound time series derived from head motion estimates and global signals were expanded with the inclusion of temporal derivatives and quadratic terms for each [75]. Frames that exceeded a threshold of 0.5 mm FD or 1.5 standardised DVARS were annotated as motion outliers. All resamplings can be performed with *a single interpolation step* by composing all the pertinent transformations (i.e., head-motion transform matrices, susceptibility distortion correction when available, and co-registrations to anatomical and output spaces). Gridded (volumetric) resamplings were performed using `antsApplyTransforms` (ANTs), configured with Lanczos interpolation to minimize the smoothing effects of other kernels [76]. Non-gridded (surface) resamplings were performed using `mri_vol2surf` (FreeSurfer).

Many internal operations of *fMRIPrep* use *Nilearn* 0.6.2 [77, RRID:SCR_001362], mostly within the functional processing workflow. For more details of the pipeline, see the section corresponding to workflows in *fMRIPrep*’s documentation.

### Analysis

#### Behavioral Analysis

We analyzed response time and accuracy performance measures for all participants, across both sessions, to ensure that participants were learning the intended visuo-motor associations. For session one, training and recall blocks in each run were analyzed separately. For session two, each run of four recall blocks was analyzed as a whole. To assess response times, we computed the mean response time for each run and participant across all correct trials. For training blocks, in line with previous work [7], [23], we excluded trials with response times > 3 SDs away from the mean for that run and participant, as well as those under 100 ms or over 3 seconds. We computed response accuracy as the fraction of trials for which participants responded correctly on the first attempt (training blocks) or correctly within two seconds (recall blocks). We then calculated the mean of these values, both within-group (modular vs. ring lattice) and across all participants, to arrive at mean values for each run (Fig. 1).

#### LS-S Trial Modeling

In our analysis, we hypothesize that the *pattern* of activation within a brain region in response to a condition contains information about that condition, beyond the information conveyed by the mean (unimodal) activation in that region. This approach, generally referred to as MVPA (Multi-Voxel Pattern Analysis), views each trial in the BOLD acquisition as a single point in an *N* -dimensional space, where *N* is the number of voxels in our region of interest, and the activity of each voxel during each trial provides coordinates in this *N* -dimensional space. Given a point for each trial, we can investigate how those points relate to one another.

Typically, BOLD timeseries are not used directly for MVPA. Instead, a typical choice for features is the resulting beta weights having fit a GLM to the data. In rapid event-based designs, such as ours, it has been suggested that running a separate GLM for each event, where all other events are represented by a single nuisance regressor (a strategy typically known as LS-S), leads to more stable estimates of beta weights [78], [79]. We used LS-S to derive a map of beta weights for each event, and for our analyses used the resulting *Z*-statistic map.

Our GLM was implemented in *nipype* using *FSL* first-level analysis routines, using the outputs (volumes and confounds) from *fMRIPrep*. All processing took place with BOLD data mapped into `MNI152NLin2009cAsym` at native resolution. Our model consisted of a double-gamma HRF plus derivatives, modeling 24 motion parameters (6-dof plus their 6 derivatives, as well as squared parameters). We used smoothing of FWHM=5mm and a high-pass cutoff of 60 seconds. Due to non-steady-state BOLD effects, we removed the first 10 TRs (or eight seconds) from each run. For each recall run, this process left us with 59 instead 60 trials, as the first trial overlapped with the non-steady-state period.

To implement LS-S, we ran 59 separate first-level analyses for each recall run—one for each trial. Each GLM consisted of the 24 motion parameters, one column for the “target” event, one column for the “nuisance” events (i.e., all remaining 58 events), and additional regressors for any motion outlier volumes as indicated by *fMRIPrep*. All columns were then convolved with the double-gamma HRF plus derivatives. We then discarded the “nusiance” regressor, and combined the 59 “target” parameter estimates across the separate first-level analyses to create our activation map across all 59 trials. These activation maps could then be used as the points in an *N* -dimensional space for our pattern-based analyses.

#### LOC Localizer

To enable robust analysis of LOC activation, participants completed a localizer run at the end of the second scanning session. This run consisted of 120 stimulus presentations and 120 phase-randomized control images, with the goal of localizing a region most sensitive to the contrast between the two. To prevent any interference from the previous learning phases, a separate set of 15 shapes were generated in the same fashion as those used in the earlier runs. These images were presented as follows: the run consisted of 30 blocks, and blocks were grouped into 10 sets of 3. Each set of 3 blocks consisted of, in order, a block of 12 stimuli, a block of 12 phase-randomized stimulus images, and a 12 second blank screen, for 36 seconds in total. Each stimulus or image was shown in the center of the screen for 0.8 s, with a 0.2 s ISI. Participants were instructed to passively view the sequence of images without responding. To control for the transition structure and the number of times each stimulus was shown, we first generated 16 connected Hamiltonian walks, for a total of 240 trials. The 240 trials were then split in half, with the first 120 assigned to object trials, and the second 120 assigned to phase-randomized trials. These 120 trials were then split between the 10 blocks, such that block 1 contained object trials 1-12 and phase-randomized trials 1-12, etc. We ran a GLM modeling shapes vs. phase-randomized trials for each subject to identify voxels that were selective to the shapes but not their phase-randomized counterparts. This GLM used the same set of nuisance regressors and same HRF convolution as in our LS-S analysis. This provided us with a *z*-scored contrast for each participant, which was used to select voxels for an LOC ROI.

#### MVPA Classification

To test whether a region coded for differences amongst trial types, we checked whether a linear classifier could predict the trial type of each recall trial based on its decoded activity pattern after being trained on independent examples of those trial types from the same subject. We used a linear SVM to predict which of the 15 trial types was most likely, and computed accuracy as the fraction of correct categorizations. Classification was performed via `LinearSVC` in *scikit-learn*. We increased the maximum number of iterations to allow convergence but otherwise used default parameters. In order to improve SVM performance, standard practice is that each feature should be zero-centered and of similar variance. Accordingly, we *z*-scored the weights for each voxel within each of the eight runs for a given participant. To reduce overfitting, we performed classification by cross-validation across runs. That is, given eight runs, we trained on seven of the eight runs, and then predicted trial identity on the remaining held-out run, and repeated for all eight runs. Cross-validated accuracy was then compared to chance performance, which here is 1/15.

#### Behavior and Classification

We hypothesized that classification accuracy might vary between modular and ring lattice graph types. Graph type has been shown to affect average response times, so we sought to test whether graph type and/or response time modulated classification accuracy. For each participant, we fit MVPA classification accuracy as a function of that participant’s mean response times on session two recall trials, in order to determine whether classification accuracy depended on response time. We fit an OLS model of classification_accuracy ∼ response_time * graph. Response time was taken as the mean response time, on correct trials (both within 2 seconds, and with the correct response), across all 480 trials comprising the eight recall runs. graph was 0 for ring lattice and 1 for modular subjects. To verify whether response accuracy predicted classification accuracy, we fit a similar model of classification_accuracy ∼ response_accuracy * graph where response accuracy ranges from 0 (missing all trials) to 1 (being perfect recall).

#### Representational Dissimilarity Matrices

We can categorize trials according to three independent attributes. First, we can use shape or, the visual appearance of the trial. Next, we can use motor movement, as in the fingers used to produce the response. Last, we can use graph location, or which other trials can precede and follow, temporally embedding the trial within either the modular or the ring lattice graph. We hypothesized that all three attributes would contribute to the neural activation evoked by a trial, and moreover contribute to how the patterns of neural activation related between trials.

For instance, we might expect that two trials share a similar representation if both responses use the ring finger. Specifically, we would predict that the trials which used ‘thumb + ring’ and ‘index + ring’ would be more similar to one another than trials which used ‘thumb + pinky’ and ‘index + ring’. Visual attributes can also impact representations—although our shapes were chosen to all be distinguishable, undoubtedly some of the shapes share features that others do not. Finally, graph structure may influence representation. If two nodes are in the same graph cluster, that cluster may be represented as well, heightening the neural similarity of those two nodes.

For a given participant, all three of these organizations are consistent—a node is always paired with both the same motor movement and the same shape—and so similarity is driven by a combination of these factors that we cannot fully disentangle. However, when we compare across participants, pairings vary: shape A might be ‘thumb’ for participant one, and ‘pinkie’ for participant two. We can therefore separate which factors drive representational similarity when we compare across participants. Comparing shapes A and B across all participants should no longer depend on the assigned motor actions, or on the location within the graph.

We asked where in the brain, and under which organization of trials, did participants have similar organizations to one another, using the concept of a noise ceiling bound. For each subject, we estimated an RDM based on a cross-validated Euclidean metric: First, within each run, we averaged LS-S beta weights for trial type, resulting in 15 activation patterns for that run. We then calculated the squared Euclidean distance between patterns, cross-validated across runs: for runs *j* and *k*, and trial types *u* and *v*, 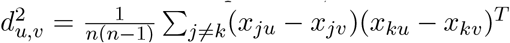, resulting in a 15 × 15 RDM for each subject. Next, we divided each matrix by its largest element, such that it ranged from 0 (all diagonal elements) to 1 (largest element), to enable cross-subject comparison. We then arranged the RDM for each subject based on the modality of comparison—either by node, motor response, or shape, such that this ordering would be consistent across subjects. Finally we took the mean of RDMs across participants separately for modular and ring lattice groups.

#### Intersubject RDM Similarity

To determine which brain regions contained similar representational structure across subjects, we conducted a whole-brain searchlight in which we correlated RDMs across subjects. First, for each subject, *PyMVPA* was used to calculate the Euclidean-distance RDM in a three-voxel radius around each voxel. The upper right triangle of the RDM was then taken, providing a 105-dimensional vector for each voxel. Next, for each subject, we correlated the RDM vector with an average vector across all other subjects, resulting in a Pearson correlation coefficient for that voxel and subject. Before comparing, RDMs were reordered to follow either graph, motor response, or shape ordering. Finally, we averaged correlation coefficients across subjects to create a map of similarity.

To statistically threshold the resulting maps, we performed a randomized null model procedure, similar to that used in Ref. [29], in order to control the familywise error rate (FWER). We repeated our RDM similarity procedure 1000 times, each time randomizing the RDM ordering by rearranging rows and columns in the RDM, separately for each subject. We then computed the mean lower bound map and found the absolute value of the largest magnitude correlation coefficient in each volume, giving us a sample distribution of 1000 correlation coefficients. We chose the 95th percentile of this sample distribution as our cutoff, and thresholded our whole-brain maps using this cutoff. Conceptually, each randomization is similar to reassigning the shape, motor, and visual correspondences for each subject. Thus, the null model procedure is identical for thresholding our similarity maps for shape, motor command, and node, and we therefore identified a single significance threshold which was used in all three maps.

#### Intrasubject RDM Consistency

Whereas the intersubject RDM similarity described above identifies shared representational organization, *intra*subject consistency can potentially identify a broader set of regions: those for which representational organization is consistent for each subject, even if the representational structure differs between subjects. Therefore, instead of comparing between subjects, we compare between runs for each subject. We employ a similar procedure to our intersubject RDM similarity: for each run, we correlate the RDM of that voxel with the mean RDM of the other seven runs, giving us a Pearson correlation coefficient for that voxel and run. We take the mean across runs to create a whole-brain map for each subject. Finally, we average those maps across subjects to identify where we observe heightened intrasubject consistency. To estimate ROI differences in intrasubject consistency, we employ a similar procedure. However, instead of operating on a whole-brain searchlight, we use our bilateral LOC ROI. Again, we calculate an RDM for each run, and then compare that RDM with the mean of the other seven runs.

#### Intrasubject Pattern Consistency

RDM similarity measures stability of relationships across runs. We may also be interested in stability of the patterns themselves. For each subject, we compute a per-run mean pattern for each of 15 trial types. We then measure the distance, using squared Euclidean distance, between that pattern and the same subject’s pattern in each of the other seven runs. We average this distance across all trial types and run combinations to arrive at a consistency value for that subject.

#### Dimensionality

To estimate dimensionality, we adapted the procedure taken in Tang, Mattar, Giusti, *et al*. [33]. We take our ability to split our data between arbitrary groupings of classes as a measure of its intrinsic dimensionality: high-dimensional data will allow more separable groupings than low-dimensional data. We begin by enumerating over both the number of classes, *m*, and permutations of *m* classes. For each value of *m*, there are 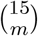 combinations of trial types, and for each combination, there are 2^*m*−1^ unique ways to divide those trial types into two groups. Dimensionality is estimated as the fraction of these assignments which we find to be linearly separable. In practice, this means that for a given value of *m*, we choose *m* out of 15 trial types, and then assign a binary label (“+” or “-”) to each of the *m* trial types. We then train an SVM to separate the “+” from the “-” trial types, and calculate the fraction of assignments above a threshold. However, in following Ref. [33], we note that, for fMRI data, this full calculation is computationally intractable and the between-subject differences can be understood from the mean classification score, rather than thresholding a classification as correct. We therefore choose a representative subsample of possible combinations and binary assignments for each value of *m*.

## Supplement

**Figure S1:**
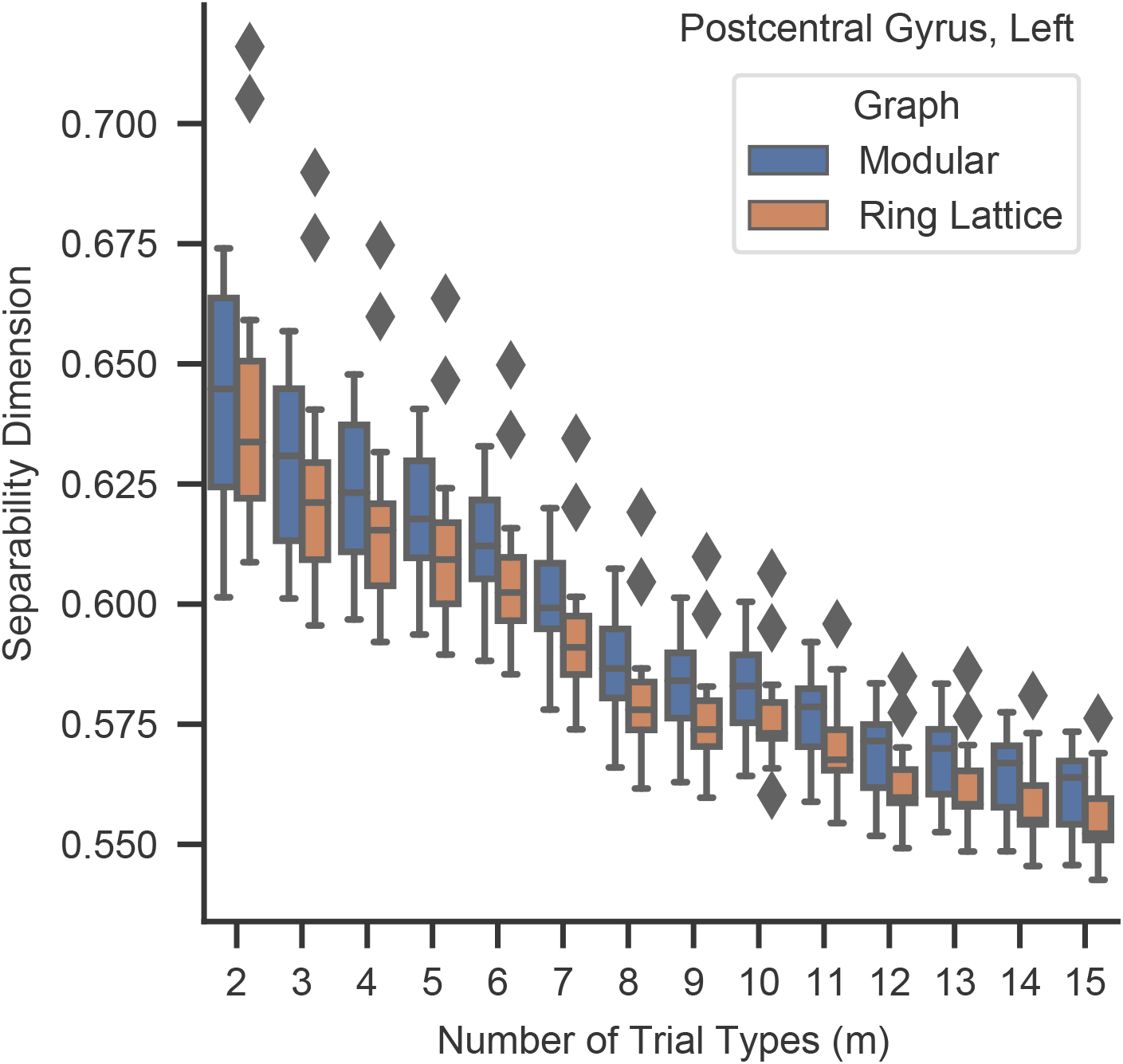
The separability dimension of the representations in postcentral gyrus. We do not observe a significant difference between graph types.

**Figure S2:**
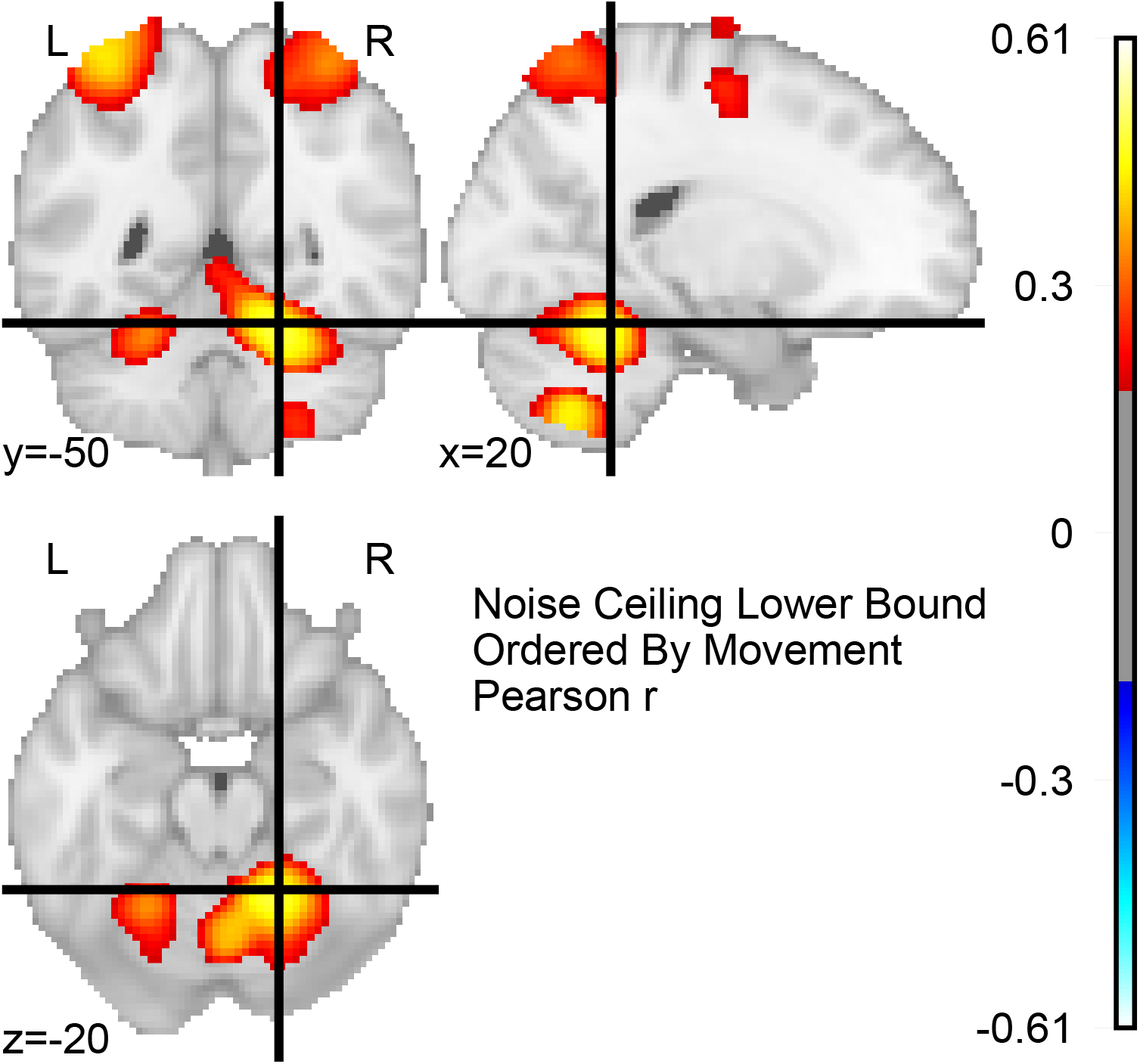
Motor RDM consistency in cerebellum. Average Pearson correlation coefficient, *r*, when the RDM in the neighborhood of a voxel is correlated with the average RDM of all other subjects in that neighborhood, centered on a region of high consistency in the cerebellum.

### Surprisal Effect

Prior work suggests that response times can reflect not only *whether* and to what degree a participant learned the graph structure, but also *how* they perceive that structure. In modular graphs, response times are longer when moving between clusters than when moving within clusters and this effect has been referred to as the cross-cluster surprisal effect [1], [2]. Although the effect is commonly observed in behavioral studies with a much larger *n* than our current study, we nevertheless examined whether cross-cluster surprisal was evident in our data. By fitting a mixed-effects linear model similar to Kahn, Karuza, Vettel, *et al*. [2], and removing trials with implausible response times (more than 3 SDs from the mean, less than 100 ms, or more than 5 s) as well as trials with incorrect responses, we estimate a surprisal effect of 24 ms (*t*_15_ = 1.80, *p <* 0.92; 95% confidence interval (CI) of -2.53 to 49.95; see Table 1). The non-significant effect is perhaps not surprising as the study was not designed to be sufficiently powered to detect it, compared with prior behavioral studies. Moreover, graph structure may modulate response accuracy alongside reaction time, and we observe a larger surprisal effect of 43 ms (*t*_15_ = 2.153, *p <* 0.048; 95% confidence interval of 3.84 to 82.16) when including trials with both correct and incorrect responses. This significant effect suggests that graph structure may similarly drive response accuracy.

**Table 1:**
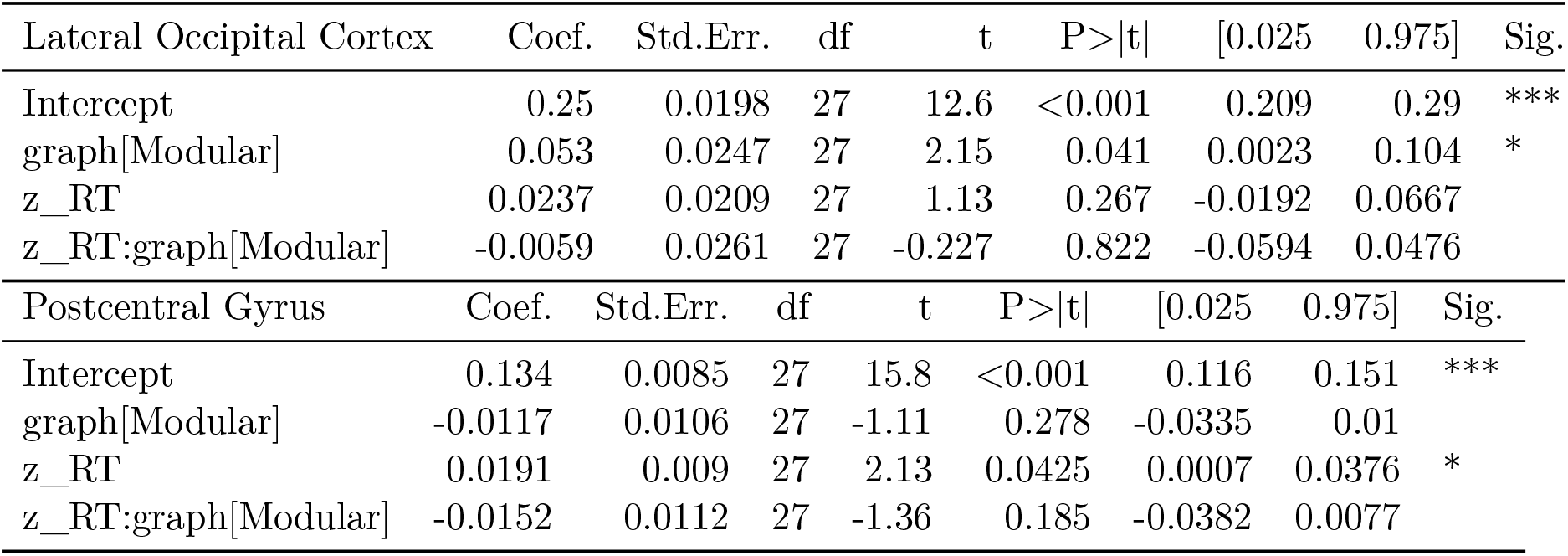
OLS regression results for classification accuracy. Both tables summarize model fit for classification_accuracy ∼ z_RT * graph where z_RT is the *z*-scored response time. Legend: +: p<0.1, *: p<0.05, **: p<0.01, ***: p < 0.001

| Lateral Occipital Cortex | Coef. | Std.Err. | df | t | P>\|t\| | [0.025 | 0.975] | Sig. |
| --- | --- | --- | --- | --- | --- | --- | --- | --- |
| Intercept | 0.25 | 0.0198 | 27 | 12.6 | <0.001 | 0.209 | 0.29 | *** |
| graph[Modular] | 0.053 | 0.0247 | 27 | 2.15 | 0.041 | 0.0023 | 0.104 | * |
| z_RT | 0.0237 | 0.0209 | 27 | 1.13 | 0.267 | -0.0192 | 0.0667 |  |
| z_RT:graph[Modular] | -0.0059 | 0.0261 | 27 | -0.227 | 0.822 | -0.0594 | 0.0476 |  |
| Postcentral Gyrus | Coef. | Std.Err. | df | t | P>\|t\| | [0.025 | 0.975] | Sig. |
| Intercept | 0.134 | 0.0085 | 27 | 15.8 | <0.001 | 0.116 | 0.151 | *** |
| graph[Modular] | -0.0117 | 0.0106 | 27 | -1.11 | 0.278 | -0.0335 | 0.01 |  |
| z_RT | 0.0191 | 0.009 | 27 | 2.13 | 0.0425 | 0.0007 | 0.0376 | * |
| z_RT:graph[Modular] | -0.0152 | 0.0112 | 27 | -1.36 | 0.185 | -0.0382 | 0.0077 |  |

To model the cross-cluster surprisal effect, we combined trials across the five training blocks for each participant who experienced the modular graph. We fit the following model using the `MixedLM` function in *stastmodels* 0.11.1 [3]:

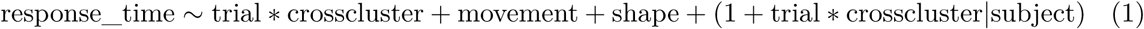

where trial is the consecutive trial number between 0 and 1500, *z*-scored to improve fitting, crosscluster is 1 at cross-cluster trials and 0 elsewhere, and both movement and shape are modeled as a categorical variables for each of the 15 possible shapes and movements. The cross-cluster surprisal effect was taken as the fixed-effect estimate of crosscluster. Our model was based on Kahn, Karuza, Vettel, *et al*. [2], with the addition of a regressor for shape. Random effects structure was chosen as the most complete such that the model converged.

To perform a power analysis for the cross-cluster surprisal effect, we conducted a bootstrap analysis using the data collected for Ref. [2]. We chose subsets of *m* participants with replacement from the full set of *n* = 30 participants used to model the cross-cluster effect in that study. We verified how often we detected a statistically significant effect, for values of *m* between 2 and 30, across 10,000 subsets for each value of *m*. We found that to achieve a false positive rate of *α* = 0.05 and power of 1 − *β* = 0.8, we would need at least 19 participants.

### Effects of Graph Learning on Visual Discrimination

How do graph-induced differences in neural representations manifest behaviorally? One possibility is that representational distance in visual regions of the brain will correspond to visual discriminatory ability when viewing those stimuli side-by-side. In the current study, neural representations of pairs of shapes in the LOC were more distinct from one another for subjects who experienced the modular graph, and repetitions of a single shape induced more consistent patterns of activation (Fig. 5). In prior work such differences in graph structure were observed to modulate the speed of subject response times when responding to a sequence of stimuli [2]: walks on modular graph structures induced lower average response times than walks on lattice graphs. Together, these results suggest that visual discrimination between stimuli might be enhanced for modular graph learners; that is, we hypothesized that learners in the current study would be able to make faster visual similarity judgements between pairs of items, assuming the neural representations induced by those items are themselves more separable.

**Table S1:** Cross-cluster surprisal mixed-effects model.

|  | Est. | Std. Err. | 95% Conf. Int. | df | t | Pr(>\|t\|) | Sig. |
| --- | --- | --- | --- | --- | --- | --- | --- |
| (Intercept) | 0.998 | 0.036 | 0.926 1.069 | 16 | 27.36 | <0.001 | *** |
| trial | -0.087 | 0.022 | -0.130 -0.044 | 15 | -3.99 | 0.0012 | ** |
| crosscluster | 0.024 | 0.013 | -0.002 0.050 | 15 | 1.80 | 0.092 | . |
| movement[01000] | -0.044 | 0.008 | -0.059 -0.029 | 20799 | -5.86 | <0.001 | *** |
| movement[00100] | -0.045 | 0.008 | -0.059 -0.030 | 20801 | -5.88 | <0.001 | *** |
| movement[00010] | 0.001 | 0.008 | -0.014 0.016 | 20774 | 0.10 | 0.92 |  |
| movement[00001] | 0.013 | 0.008 | -0.002 0.029 | 20770 | 1.66 | 0.097 | . |
| movement[11000] | 0.123 | 0.008 | 0.108 0.138 | 20819 | 15.76 | <0.001 | *** |
| movement[01100] | 0.078 | 0.008 | 0.063 0.094 | 20801 | 9.78 | <0.001 | *** |
| movement[00110] | 0.133 | 0.008 | 0.118 0.148 | 20775 | 17.30 | <0.001 | *** |
| movement[00011] | 0.176 | 0.008 | 0.161 0.192 | 20788 | 22.78 | <0.001 | *** |
| movement[10100] | 0.227 | 0.008 | 0.212 0.243 | 20798 | 29.13 | <0.001 | *** |
| movement[01010] | 0.183 | 0.008 | 0.167 0.198 | 20779 | 23.58 | <0.001 | *** |
| movement[00101] | 0.365 | 0.008 | 0.349 0.381 | 20826 | 44.62 | 0 | *** |
| movement[10010] | 0.201 | 0.008 | 0.186 0.216 | 20807 | 25.92 | <0.001 | *** |
| movement[01001] | 0.195 | 0.008 | 0.180 0.210 | 20794 | 24.86 | <0.001 | *** |
| movement[10001] | 0.150 | 0.008 | 0.134 0.166 | 20805 | 18.72 | <0.001 | *** |
| shape2 | -0.044 | 0.008 | -0.059 -0.029 | 20816 | -5.76 | <0.001 | *** |
| shape3 | -0.013 | 0.008 | -0.028 0.002 | 20804 | -1.70 | 0.089 | . |
| shape4 | -0.107 | 0.008 | -0.122 -0.092 | 20735 | -14.04 | <0.001 | *** |
| shape5 | -0.067 | 0.008 | -0.082 -0.053 | 20766 | -8.84 | <0.001 | *** |
| shape6 | -0.077 | 0.008 | -0.092 -0.062 | 20819 | -10.22 | <0.001 | *** |
| shape7 | -0.027 | 0.008 | -0.042 -0.012 | 20825 | -3.49 | <0.001 | *** |
| shape8 | -0.028 | 0.007 | -0.042 -0.013 | 20811 | -3.74 | <0.001 | *** |
| shape9 | -0.081 | 0.008 | -0.095 -0.066 | 20822 | -10.74 | <0.001 | *** |
| shape10 | -0.126 | 0.008 | -0.141 -0.111 | 20722 | -16.34 | <0.001 | *** |
| shape11 | -0.073 | 0.008 | -0.088 -0.058 | 20790 | -9.47 | <0.001 | *** |
| shape12 | -0.047 | 0.008 | -0.061 -0.032 | 20782 | -6.13 | <0.001 | *** |
| shape13 | -0.043 | 0.008 | -0.059 -0.028 | 20818 | -5.61 | <0.001 | *** |
| shape14 | -0.023 | 0.008 | -0.038 -0.008 | 20821 | -3.05 | 0.0023 | ** |
| shape15 | -0.134 | 0.008 | -0.149 -0.119 | 20778 | -17.48 | <0.001 | *** |
| trial:crosscluster | 0.008 | 0.009 | -0.010 0.027 | 20819 | 0.91 | 0.36 |  |

To test this hypothesis, we recruited an independent group of subjects to perform binary similarity judgements on stimulus pairs after training on either the modular or lattice graph. Using the same set of stimuli as in our fMRI group, we exposed each participant to a walk on either a modular or lattice graph. We then displayed pairs of stimuli to the subject, either distinct or identical, instructed subjects to respond as quickly as possible with one key if the shapes were distinct, another if the shapes were identical, and measured their resulting response times as an indicator of visual discriminability. We found that response times for pairs of shapes are highly correlated with the neural dissimilarity of those shapes: when presented with shapes whose LOC activation patterns were more dissimilar from one another, subjects responded more quickly than when given pairs whose LOC activation patterns were more similar. However, response times did not significantly differ between the modular and lattice graph conditions. Further details of this additional study can be found below.

### Participants

Fifty two participants (19 male, 29 female, two who declined to report gender, plus two participants later excluded) were recruited from the general population of Philadelphia, Pennsylvania. All participants gave written informed consent and the Institutional Review Board at the University of Pennsylvania approved all procedures. In line with the neuroimaging subject pool, all subjects were right-handed, between the ages of 18 and 35 years (M=24.1; SD=4.32), and had no known psychiatric disorders or past neurological medical history.

### Subject Preparation

The experimenter situated the subject in front of the laptop and began the experiment, which included instructions on each day for performing the task. The experimenter only clarified what was mentioned in the on-screen instructions. Subjects wore headphones; during session one this was only to block out sound, but session two included auditory cues for the task.

### Stimuli and Responses

Subjects were trained and tested on a set of 15 visual stimuli, and 15 one- or two-key combinations, using the keys ‘space’, ‘j’, ‘k’, ‘l’, and ‘;’. Stimuli were generated in MATLAB using the ShapeToolbox, by perturbing a sphere with sinusoids (see earlier section on fMRI Methods for details). As before, each of the 15 shapes and motor responses was assigned to one of 15 nodes in a graph. Each mapping was random and unique to that participant. There were two possible graphs: a ring lattice where 15 nodes are arranged in a ring and each is connected to its two nearest neighbors in each direction, and a modular graph, where 15 nodes are arranged into three densely connected clusters, with one edge connecting each pair of clusters.

### Session One Design

In session one, participants completed 5 runs of 300 trials, identical to session one for the fMRI participants, only differing in that the five square outlines corresponded to five keys on a keyboard (‘space’, ‘j’, ‘k’, ‘l’, and ‘;’) rather than buttons on an MRI-compatible button box. As before, trial order was generated based on a walk through the graph; in other words, if the stimulus (and associated motor response) of the current trial correspond to node X in the graph, then the stimulus (and associated motor response) of the next trial correspond to one of the neighbors of X, continuing in an unbroken sequence for 300 trials.

### Session Two Design

In session two, participants completed 8 runs of 210 trials of a binary classification task on pairs of presented shapes, rather than reproducing the button combination of each shape as was done in session two for the fMRI subjects. On each trial, 2 of the 15 stimuli were shown side-by-side. The stimuli were either both images of the same shape, differing in only angle and size, or 2 entirely different shapes, and participants were instructed to indicate as quickly as possible whether the images were different shapes (by pressing ‘f’) or the same shape (by pressing ‘j’). Rather than testing on all 105 unique shape pairs, each subject was tested on 1 of 5 predetermined sets of 21 shape pairs to enable more sampling of each pair within-subject. Combining the 5 sets across subjects covered the space of all shape pairs. Stimulus onset timing was jittered, varying from 800 ms to 1.3 s after the start of the trial. Stimuli remained on the screen for 500 ms until being replaced by a fixation target. Subjects had an additional 1.5 s to respond after the stimuli disappeared. Subjects were presented with auditory cues to both indicate trial timing and errors: a 440Hz beep indicated the start of every stimulus display, a 300Hz beep was played to indicate an incorrect response, and a 250Hz beep was played to indicate that the subject failed to respond in time.

## Results

For each subject, we calculated the mean inverse response time when responding to each pair of stimuli, excluding incorrect responses, as a measure of visual dissimilarity.

### Behavioral Similarity Judgements and Neural Dissimilarity Measurements

Here we ask whether the inverse response time to each pair of stimuli was correlated with the LOC representational dissimilarity score between those same stimuli. That is, whether pairs of stimuli which elicited faster response times in the behavioral data were also those eliciting more dissimilar neural representations in the neuroimaging data. Since stimulus location is randomized in graph location across both experiments, here we are solely investigating effects due to visual properties of the stimuli. To test this relationship, we took the mean inverse response time of all similarity judgements across subjects in the behavioral data, and calculated Pearson’s correlation coefficient, *r*, with the average *z*-scored RDM value across all subjects in the neuroimaging data for that same pair of shapes. We found a significant correlation between behavioral similarity judgements and the neural dissimilarity (Fig. S3, Pearson’s *r* = 0.3, *t*_103_ = 3.3, *p <* 0.002), suggesting that visual properties of the stimuli significantly drive both measures.

### Effect of Graph Structure on Discrimination Judgements

Next, we asked whether the differences observed in neural representations driven by graph type were likewise reflected in visual discriminability judgements. Amongst our neuroimaging dataset subjects, we found that exposure to the modular graph induced more distinct representations in LOC than did exposure to the lattice graph. Given that visual discriminability judgements were strongly correlated with neural dissimilarity, we hypothesized that exposure to the modular graph would enable more rapid visual discrimination between the stimuli than exposure to the lattice graph.

To investigate the effect of graph structure on discrimination judgements, we fit a linear mixed effects model to the response times collected on the second day of the behavioral experiment (1/RT ∼ Graph * LOC Dissimilarity + (1 | Subject) + (1 | Shape Pair)). We predicted inverse RTs based on the graph type experienced by the subject (Lattice / Modular) and the LOC dissimilarity score for that pair of shapes, and included random effects both for subject and for the shape pair. We excluded incorrect responses as well as match trials where both stimuli were the same. Dissimilarity was taken as the mean RDM value for that pair of shapes across both lattice and modular fMRI groups, then *z*-scored across all shape pairs, in order to account for graph-independent variance due to the visual properties of stimuli.

Here we report fixed effects from the model (Table S2). We find a significant effect of LOC dissimilarity on performance (*t*_93_ = 4.33, *p <* .0001), in line with the high correlation previously observed between LOC dissimilarity and response times. However, when we asked whether graph exposure differentially affected performance between the two groups, we found no significant differences between lattice and modular graphs (*t*_48_ = 0.866, *p <* 0.392).

**Figure S3:**
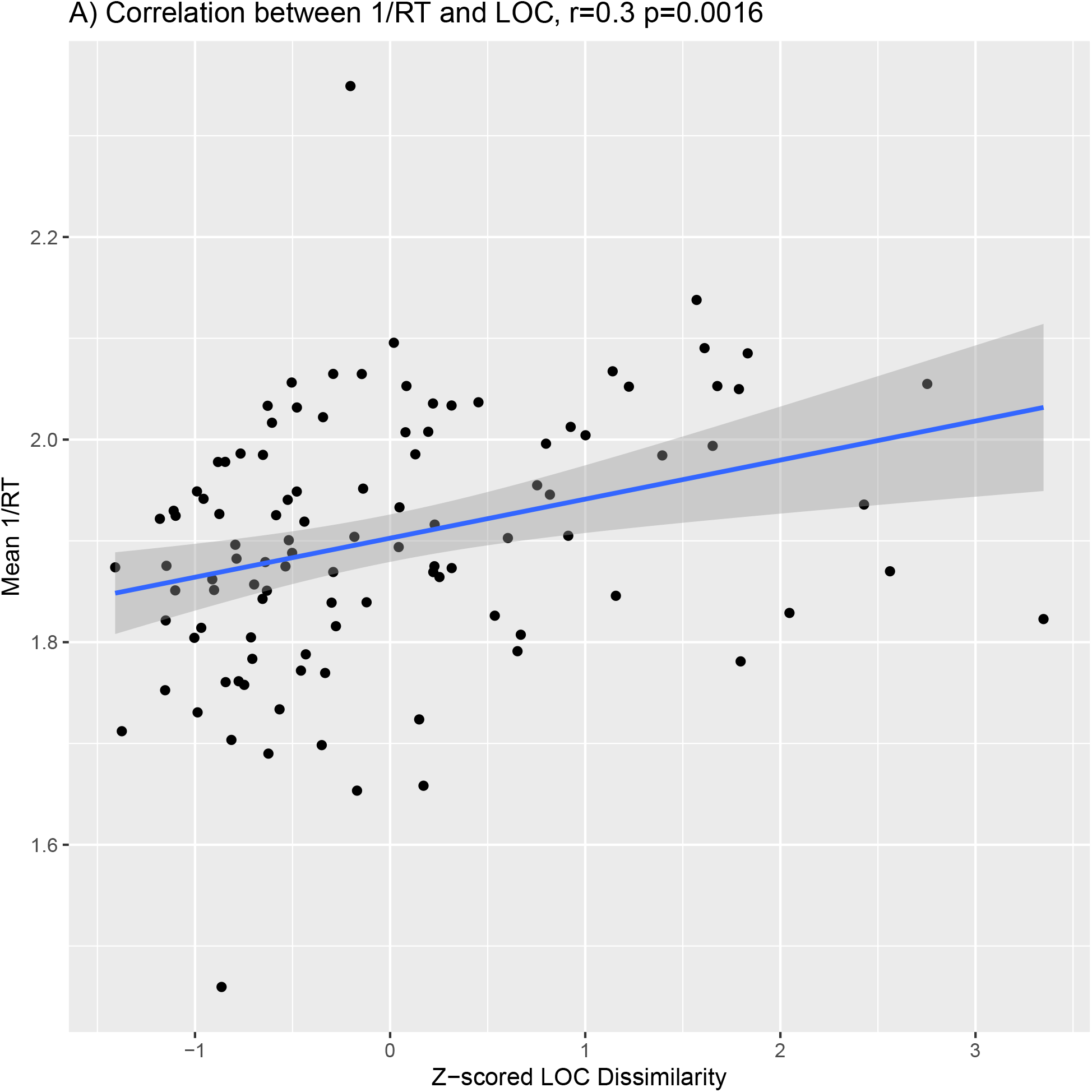
Discrimination judgement inverse response times are correlated with LOC neural dissimilarity. Significant correlation between mean inverse response time averaged across all subjects for each shape pair and the corresponding *z*-scored LOC dissimilarity, averaged across all subjects in both modular and lattice conditions (Pearson’s *r* = 0.3, *t*_103_ = 3.3, *p <* 0.002). Each point corresponds to a stimulus pair.

**Table S2:** Graph Difference Fixed Effects. We fit a mixed effects model to data from the behavioral experiment, and predicted the inverse RT on each non-match trial based on graph type the subject was exposed to (Lattice / Modular) and an LOC dissimilarity score for the tested shape pair, with random effects for both subject and shape pair. Dissimilarity was taken as the mean RDM value for that pair of shapes across both lattice and modular fMRI groups, then *z*-scored across all shape pairs. Here we report fixed effects from the model. We find a significant effect of LOC dissimilarity on performance. We find no significant differences between lattice and modular graphs. Significance codes: 0 ‘***’ 0.001 ‘**’ 0.01 ‘*’ 0.05 ‘.’ 0.1 ‘ ‘ 1.

| Lattice vs. Modular | Estimate | df | t value | Pr(>\|t\|) |  |
| --- | --- | --- | --- | --- | --- |
| (Intercept) | 1.871 | 48.809 | 34.496 | < 2e-16 | *** |
| Graph (Lattice) | 0.066 | 47.704 | 0.866 | 0.391 |  |
| LOC RDM value | 0.034 | 93.179 | 4.332 | 3.72e-05 | *** |

